# Conserved spatial patterning of gene expression in independent lineages of C_4_ plants

**DOI:** 10.1101/2025.05.14.654047

**Authors:** Tianshu Sun, Venkata Suresh Bonthala, Benjamin Stich, Leonie Luginbuehl, Julian M. Hibberd

## Abstract

- C_4_ photosynthesis enhances carbon fixation efficiency by reducing photorespiration through the use of an oxygen-insensitive carboxylase and spatial separation of photosynthesis between mesophyll and bundle sheath cells. The C_4_ pathway has evolved independently in more than sixty plant lineages but molecular mechanisms underpinning this convergence remain unclear. To explore this, we generated high-resolution transcriptome atlases for two independently evolved C_4_ dicotyledonous species - *Gynandropsis gynandra* (NAD-ME subtype) and *Flaveria bidentis* (NADP-ME subtype).
- We used both single-cell and single-nucleus RNA sequencing to capture gene expression profiles from individual leaf cells, enabling detailed comparison of cell types and transcriptional signatures.
- Both approaches produced biologically comparable data for major leaf cell types, transcriptomes from single-nucleus sequencing showed lower stress signatures and were more representative of tissue proportions in the leaf. The single-nucleus data revealed that bundle sheath cells from both C_4_ species share a gene expression pattern associated with mesophyll cells of C_3_ plants. A conserved set of transcription factors, including members of the C2H2 and DOF families, was identified in the bundle sheath cells of both species.
- This study presents the first single-cell-resolution transcriptomes for two independent C_4_ dicot lineages and provides a valuable resource, including a web-based portal for data visualization.

## Introduction

Photosynthesis sustains much of life on Earth. Central to the photosynthetic process is the enzyme Ribulose-1,5-Bisphosphate Carboxylase-Oxygenase (RuBisCO), which fixes atmospheric carbon dioxide into the C_3_ compound 3-phosphoglyceric acid. While RuBisCO predominantly catalyses this carboxylation reaction, it can also bind and react with oxygen to produce the toxic metabolite 2-phosphoglycolate. The photorespiratory pathway allows the metabolism of 2-phosphoglycolate, but requires additional energy input and leads to the loss of fixed nitrogen. Because the rate of oxygenation of RuBisCO increases with temperature, photorespiration becomes increasingly significant close to the equator (Schluter & Weber, 2020; Smith et al., 2023). To mitigate these effects, multiple lineages of plants have evolved carbon-concentrating mechanisms that reduce the oxygenation reaction of RuBisCO, and thus rates of photorespiration. One such example is C_4_ photosynthesis (Hibberd & Covshoff, 2010; Ben P. Williams, Aubry, & Hibberd, 2012).

In contrast to the majority of plants where photosynthesis is primarily associated with mesophyll cells, in C_4_ species the process is typically equally divided between cell types such as the mesophyll and bundle sheath. Thus, in C_4_ plants, carbon fixation operates across different cell types rather than within a single cell. Initially, HCO_3_^−^ is fixed in the mesophyll by phospho*enol*pyruvate carboxylase (PEPC) into C_4_ acids, which can diffuse to the bundle sheath. Subsequent decarboxylation of these C_4_ acids in the bundle sheath creates a CO_2_-rich environment around RuBisCO. This compartmentation of the photosynthetic process enhances carbon fixation efficiency and reduces the need for photorespiration. However, the transition from C_3_ photosynthesis to the two-celled C_4_ pathway requires significant optimisation of leaf anatomy, cell biology and biochemistry - all of which are underpinned by the rewiring of pre-existing gene networks (Schluter & Weber, 2016). Despite this complexity, C_4_ photosynthesis has independently evolved more than 60 times across multiple lineages of land plants (R. F. Sage, Christin, & Edwards, 2011). Although some C_4_ traits are underpinned by parallel evolution (Aubry, Kelly, Kumpers, Smith-Unna, & Hibberd, 2014; Brown et al., 2011; Christin, Salamin, Savolainen, Duvall, & Besnard, 2007), it has been known for some time that many characteristics of C_4_ photosynthesis are convergent (Christin & Osborne, 2013; R. F. Sage, Sage, & Kocacinar, 2012; B. P. Williams, Johnston, Covshoff, & Hibberd, 2013). This includes for example variation in Kranz anatomy as well as the biochemical basis of the pathway. For example, whilst in most cases bundle sheath cells have been co-opted for the photosynthetic carbon reduction cycle, in some species use the mestome sheath for this purpose (Khoshravesh et al., 2020; Khoshravesh et al., 2016; R. F. Sage, 2004). Moreover, enzymes used to decarboxylate C_4_ acids during the photosynthetic carbon reduction cycle belong to three classes of protein - NADP-malic enzyme (NADP-ME), NAD-malic enzyme (NAD-ME), and PEP carboxykinase (PEPCK) (Niklaus & Kelly, 2019; Schluter & Weber, 2020). The molecular basis of this convergent evolution of C_4_ traits however remains poorly understood (Westhoff & Gowik, 2010).

One reason for limited progress in understanding the molecular basis of C_4_ photosynthesis is associated with the complexity of separating specific cell types from leaves. Mechanical separation of cell types allowed mesophyll cells to be isolated from bundle sheath strands, but the latter are composed of xylem and phloem as well as the bundle sheath itself (Covshoff, Furbank, Leegood, & Hibberd, 2013; Sheen, 1995). Protoplasting has also been used to isolate and study mesophyll and bundle sheath strands, but the process of isolating protoplasts induces stress responses to gene expression (Sawers, Liu, Anufrikova, Hwang, & Brutnell, 2007). We reasoned that the advent of single-cell and single-nucleus sequencing technologies may provide tools to better understand the molecular basis of the convergent evolution of C_4_ photosynthesis. For example, droplet-based single-cell RNA-SEQ technologies have significantly increased the throughput and resolution of transcriptome analyses (Chen, Ge, & Lu, 2023; Cuperus, 2022). However, capturing the full diversity of cells in complex tissues using single-protoplast RNA-SEQ can be challenging due to the trade-off between recovery of all cell types and longer incubation periods required for release of these cells causing stress responses in gene expression (Kim et al., 2021). In leaves, this issue can be exacerbated where vascular and bundle sheath cells are located deep in the tissue and digestion of cell walls is slow. Moreover, even after optimising protoplasting, some cell types can remain under-represented, with for example less than 2% bundle sheath cells being captured, and limited numbers of vascular or epidermal being retained (Bezrutczyk et al., 2021). In mammalian studies, single-nucleus RNA-SEQ has proved valuable for analyzing tissues, such as neurons (Lacar et al., 2016), which are challenging to dissociate into single-cell suspensions. It has also helped minimize gene expression changes that may arise from dissociation (Ding et al., 2020; van den Brink et al., 2017). Although transcript abundance is lower than in single-cell RNA-SEQ, the single-nucleus method has demonstrated strong sensitivity and effectively classified cell types (Bakken et al., 2018; Guillotin et al., 2023; Lake et al., 2017). Moreover, recently single nucleus RNA-SEQ has been used to assess gene expression during photomorphogenesis in C_3_ and C_4_ grasses (Swift, Luginbuehl et al 2024).

We therefore set out to compare single-cell and single-nucleus RNA-sequencing from two species that evolved C_4_ photosynthesis independently. Specifically, we chose *Gynandropsis gynandra* and *Flaveria bidentis* that belong to the dicotyledons, have previously been used to study C_4_ photosynthesis, and have annotated genome sequences. *G. gynandra* (formerly Cleome) uses NAD-ME as the primary decarboxylase in bundle sheath cells (Brown, Parsley, & Hibberd, 2005; Marshall et al., 2007), is a leafy vegetable widely grown across Sub-Saharan Africa and South East Asia (Mashamaite, Manyevere, & Chakauya, 2022), and is sister to the Brassicaceae and therefore closely related to *Arabidopsis thaliana*. Comparative analysis between *Gynandropsis gynandra* and *A. thaliana* has provided insight into multiple aspects of the complex C_4_ phenotype including the formation of plasmodesmata (Schreier et al., 2024), evolution of the genome (Hoang et al., 2023), and the rewiring of gene networks responsive to light, as well as associated with cell specific gene expression and cell identity (Singh et al., 2023). *F. bidentis* uses NADP-ME as the primary C_4_ acid decarboxylase (Gowik, Brautigam, Weber, Weber, & Westhoff, 2011), and comparative analysis of species in the genus formed the foundations for many of the conceptual and mathematical models that are used to explain why and how C_4_ photosynthesis evolved (Heckmann et al., 2013; T. L. Sage et al., 2013; Schulze et al., 2013). *G. gynandra* and *F. bidentis* belong to the rosid and asterid angiosperm clades, the two largest eudicot clades, which diverged at least 100 million years ago (Moore, Soltis, Bell, Burleigh, & Soltis, 2010; Zuntini et al., 2024).

We compiled single-protoplast and single-nucleus transcriptome atlases for all major cell types of *G. gynandra* and *F. bidentis* leaves, and provide a browser-based interactive tool built with ShinyCell to allow exploration of the data. Both protoplast and nucleus sequencing produced biologically meaningful and comparable cell-type-specific data for the major cell types in a C_4_ leaf, but single nuclei provided more accurate cell type representation, and transcriptomes lacked protoplast-induced stress responses. Analysis of transcript abundance in all major cell types of leaves allowed us to investigate the extent to which mesophyll and bundle sheath cells from these independent C_4_ lineages shared the same molecular signatures. Despite representing distinct C_4_ lineages and using different decarboxylation enzymes in the bundle sheath, in both *G. gynandra* and *F. bidentis* compared with the ancestral C_3_ state, bundle sheath cells gained expression of a similar set of genes, previously associated with mesophyll cells and allowing decarboxylation and carbon fixation in the C_4_ cycle. Furthermore, we identify transcription factor families, such as IDD and DOFS, that are preferentially expressed in either mesophyll or bundle sheath cells of both species.

## Materials and Methods

### Plant growth conditions

*Gynandropsis gynandra* seeds were germinated on moist filter paper in the dark at 32°C for 32 hours, then sown directly onto soil. *Flaveria bidentis* seeds were placed on ½-strength Murashige and Skoog medium in 0.8% (w/v) agar (pH 5.8) and grown for 10 days in a growth chamber at 24°C, photoperiod 16 hours light and 8 hours dark before being transferred to soil. After sowing *G. gynandra* and transplanting *F. bidentis*, plants were maintained in a growth cabinet under a 16-hour light/8-hour dark cycle at 24°C, with 60% relative humidity, and a photon flux density of 400 μmol m^−2^ s^−1^. Plants were grown for 6 weeks until leaves were harvested for single-protoplast or single-nucleus RNA sequencing. To generate DNA for genome assembly, *F. bidentis* seeds were germinated and grown in growth chambers at Heinrich Heine University, Dusseldorf, Germany. Samples from multiple organs – leaves, stem and root were collected and frozen in liquid nitrogen. Following the manufacturer’s protocol, genomic DNA was extracted from leaves and RNA from all organs above-mentioned using the Qiagen DNEasy Mini Kit (Qiagen, Germany).

### Whole-genome sequencing, estimation of genome size, reference-quality genome assembly and genome annotation

The genome of *Flaveria bidentis* was sequenced at NRGene, Nes Ziona, Israel using 10X linked-read and Illumina sequencing technologies. Genome size was estimated using the K-mer (127 bp) coverage distribution and the heterozygous (Het) and homozygous (Homo) peaks of unique K-mer distributions were chosen for genome size estimation. The estimated genome size was then calculated by the division of all K-mer by the coverage of unique K-mers of Het peaks.

The 10X linked-read and Illumina sequence libraries were used for *de novo* genome assembly using the DeNovoMAGIC pipeline (NRGene, Nes Ziona, Isreal), a DeBruijn-graph-based genome assembler to generate a reference-quality assembly. We assessed the contiguity of the assembly using the N50 metric. The completeness of the assembly was assessed by evaluating the presence of conserved single-copy orthologous genes using Benchmarking Universal Single-Copy Orthologs (BUSCO) v5.5.0 against the “embryophyta_odb10” lineage dataset (Seppey et al., 2019).

We performed gene annotation using Helixer v0.3 (Stiehler et al., 2021), a deep learning-based gene annotation tool with a training dataset of “land_plant” to annotate protein-coding genes. Further, we performed functional annotation for predicted genes using the standalone version of InterProScan v5.57-90.0 (Jones et al., 2014). In addition, gene descriptions were assigned by performing BLASTP (Altschul et al., 1990) with default parameters against the UniProt protein database (downloaded on 29/August/2022 from https://www.uniprot.org). Finally, gene models were removed which produced protein sequences with <50 amino acids and gene models related to transposable elements using a curated list of Pfam protein domain IDs (Jayakodi et al., 2020).

We combined the extracted RNA from multiple organs into a single library and sequenced it using Oxford Nanopore direct RNA sequencing (DRS) technology at the Genomics & Transcriptomics Lab, Heinrich Heine University, Düsseldorf, Germany. The DRS reads were subjected to quality filtering using NanoFilt v2.8.0 (De Coster et al., 2018) to remove reads with a length <150 bases and quality q < 10, followed by trimming adapters using Porechop v0.2.4 (Wick et al., 2017). Finally, the high-quality reads were mapped against the reference genome and expression abundance (TPM) was estimated using the NanoCount v1 (Gleeson et al., 2022). Genes with TPM > 0 were considered as expressed.

### Protoplast and nuclei isolation

For protoplast isolation, leaves were harvested and finely chopped in a 9cm petri dish containing 15 mL of freshly prepared enzyme cocktail (1.25% (w/v) cellulase RS, 0.3% (w/v) macerozyme R10, 0.2–0.6 M mannitol, 20 mM KCl, 20 mM MES, pH 5.7, 10 mM CaCl₂, and 0.1% (w/v) BSA). Tissue was incubated for 2.5 hours on a shaking incubator (40 rpm, 25°C, dark) and digestion stopped by adding an equal volume of W5 buffer (2 mM MES, pH 5.7, 125 mM CaCl₂, 154 mM NaCl, and 5 mM KCl). The released protoplasts were filtered twice using a 40 µm cell strainer then collected in a 50 mL tube. Protoplasts were centrifuged at 100 x *g* for 5 minutes and resuspended three times — twice in W5 buffer and then 0.4 M mannitol buffer for purification. After the final wash, the supernatant was gently removed, leaving 0.5–1 mL of protoplast suspension in 0.4 M mannitol. Protoplast integrity and concentration were assessed using a C-Chip hemocytometer under a light microscope. Protoplast suspensions of good quality (>80% integrity, minimal debris) were adjusted to a concentration of approximately 1000 protoplasts/µL.

Nuclei were isolated according to the published protocol (Swift et al., 2024). Briefly, leaves were harvested at the same development stage and finely chopped in a 5 cm Petri dish containing 1.5 mL of freshly prepared nuclei purification buffer (10 mM Tris-HCl, pH 7.4, 10 mM NaCl, 3 mM MgCl₂, 0.5 mM spermidine, 0.2 mM spermine, 1X Roche Complete protease inhibitors, 0.2% (w/v) BSA, and 1 U/µL Protector RNase Inhibitor). Tissue was filtered once through a 70 µm and then through a 40 µm cell strainer. Nuclei were stained with Hoechst and sorted via FACS. Sorted nuclei were collected in a 1.5-mL tube containing BSA and Protector RNase Inhibitor, then counted using a C-Chip hemocytometer under a light microscope. Nuclei suspensions were adjusted to a concentration of approximately 400–600 nuclei/µL.

### Library preparation and sequencing, data processing and quality control

A total of 10,000 protoplasts (targeting 6,000 recovered cells) or 16,000 nuclei (targeting 10,000 recovered nuclei) were loaded per well according to the Chromium Next GEM Single Cell 3ʹ Reagent Kits v3.1 protocol. Following the manufacturer’s standard protocol, cDNA for each sample was barcoded, amplified, and single-cell libraries were constructed (dual index), then sequenced using Illumina NovaSeq systems.

Demultiplexed sequences were analysed using CellRanger v7.0.0 (10x Genomics) with default parameters to generate initial gene–cell matrices (using intron mode). All samples were processed with default settings, except for the nuclei libraries from *G. gynandra*, which were analysed with -- force-cells = 10,000. Reads from *G. gynandra* were mapped to its genome assembly (v3.0) (Hoang et al., 2023), and reads from *F. bidentis* mapped to the reference-quality genome assembly that we generated in this study. To account for ambient RNA contamination, count matrices were adjusted using the adjustCounts() function from SoupX (Young & Behjati, 2020). DoubletFinder (McGinnis, Murrow, & Gartner, 2019) was used to assess the probability of droplets containing more than one nucleus or protoplast, and predicted doublets were excluded from downstream analysis. Further quality control was performed using Seurat v4.3.0 (Hao et al., 2021). Genes expressed in fewer than three cells, and cells with fewer than 200–500 genes (depending on dataset quality) were excluded from further analyses.

### Clustering, cell type annotation, homolog identification, gene ontology and orthology analysis

Nuclei or protoplast libraries from the same species were merged using Seurat v4.3.0 and batch-corrected with harmony v0.1.1 (Korsunsky et al., 2019). For integration of nuclei and protoplast libraries, Seurat’s FindIntegrationAnchors() function was used to identify anchor genes, followed by integration using IntegrateData(). Neighbor graph construction, cluster determination, and nonlinear dimensionality reduction were performed with Seurat using FindNeighbors(), FindClusters(), and RunUMAP() functions.

Homologs between *Arabidopsis thaliana, G. gynandra* and *F. bidentis* were identified using blastp v2.12.0 (Camacho et al., 2009). Nuclei or protoplast clusters were annotated based on the expression of homologous genes corresponding to known leaf cell type markers from Arabidopsis. Clusters without clear marker gene expression or showing mixed expression patterns were labelled as “ua” (unable to annotate). Annotation information was added to nuclei and protoplasts in the Shiny-based web application for atlas visualization, which was created using ShinyCell v2.1.0 (Ouyang, Kamaraj, Cao, & Rackham, 2021). Marker genes for each cell type were identified using Seurat’s FindAllMarkers() function by comparing nuclei/protoplasts from a specific cell type against all other cell types. Differentially expressed genes (DEGs) between nuclei and protoplasts, or between two different cell types, were identified using FindMarkers(). Marker genes and DEGs were only considered if they showed a log fold change of at least 0.25 and were detected in a minimum fraction of 10% of cells in either population. The best BLAST hit against Arabidopsis of each marker genes/DEGs were used for Gene ontology (GO) enrichment analyses, and performed using clusterProfiler v4.2.2 (Wu et al., 2021), with a false discovery rate (FDR) cut-off of 0.05. Stress and nuclei signature scores were calculated with the AddModuleScore() function in Seurat using the expression of genes enriched in respective pathways.

Compartmentation of orthologous genes across cell types in *A. thaliana, G. gynandra* and *F. bidentis* was based on orthogroups identified by OrthoFinder v2.4 (Emms & Kelly, 2019, using protein sequences from primary transcripts of species including *Zea mays* (RefGen_V4), *Oryza sativa* (v7.0), *Brachypodium distachyon* (v3.2), *Sorghum bicolor* (v3.1), *Vitis vinifera* (v2.1), *Theobroma cacao* (v2.1), *A. thaliana* (Araport11), *Carica papaya* (ASGPBv0.4), *Malus domestica* (v1.1), *Fragaria vesca* (v4.0.a2), *Ricinus communis* (v0.1), *Manihot esculenta* (v8.1), *Populus trichocarpa* (v4.1), *Medicago truncatula* (Mt4.0v1), *Glycine max* (Wm82.a6.v1), *Solanum lycopersicum* (ITAG5.0), and *G. gynandra* and *F. bidentis*. The Arabidopsis single-protoplast expression matrix (Kim et al., 2021) was used to compare cell type expression patterns between C_3_ and C_4_ plants. Transcription factor families were classified based on the PlantTFDB (Jin et al., 2017).

## Results

### Optimised protocols for single nucleus and protoplast isolation from *Gynandropsis gynandra* and *Flaveria bidentis*

To allow a direct comparison of single nucleus and single protoplast RNA-seq from leaves of C_4_ plants, we optimised experimental protocols to obtain high-quality suspensions of nuclei and protoplasts from *Gynandropsis gynandra* and *Flaveria bidentis* leaves (Fig. S1). Nuclei were isolated after chopping leaves, and individual nuclei then purified using fluorescence-activated cell sorting. To ensure clear separation of nuclei from debris and chloroplasts, we adjusted the gating for positive events by excluding those with high autofluorescence (Fig. S2). Purified nuclei were then used for library preparation. For protoplast isolation, enzyme composition, mannitol concentration, and age of plants were assessed. Under our conditions, age had a major effect on protoplast viability – with those from older plants being less likely to burst (Fig. S3a,b). We also found that avoiding the commonly used protoplast purification solutions, which contain Na⁺, K⁺ and Ca²⁺, was crucial for efficient transcriptome capture, as its presence ultimately resulted in decreased gene detection (Fig. S3c-f).

With these optimised samples we generated libraries using the 10x Chromium platform from nuclei and protoplasts of *G. gynandra* and *F. bidentis* and subjected these to sequencing. Reads from *Gynandropsis gynandra* were mapped to the published genome (Hoang et al., 2023), while reads from *Flaveria bidentis* were mapped to a reference-quality genome we assembled for *Flaveria bidentis* with N50 of 41.8 Mb capturing 96.29% of complete BUSCO genes of the embryophytes (Table S1) (Seppey et al., 2019). To generate this genome we sequenced ∼884 Gb that represented a coverage of approximately x635 (Table S2). The genome size of *Flaveria bidentis* as 1.39 Gb was estimated with the K-mer coverage distribution approach, and this led to a predicted a greater size (1.01 Gb) than that previously reported from flow-cytometry (Taniguchi et al., 2021). Genome annotation produced 26,603 gene models with an average length of 3692.35 bases, and functional annotation assigned protein domain descriptions (PFAM) and gene ontology (GO) terms to 80.4% and 52.79% of genes respectively.

After mapping raw reads to the respective genomes and performing doublet removal, replicate combination, quality control and batch effect removal, we generated gene expression matrices for all four datasets. For *G. gynandra*, we identified 22,224 genes from 18,043 nuclei (913 genes per nucleus) and 23,522 genes from 5,758 protoplasts (3,974 genes per protoplast) (Fig. 1a-c). For *F. bidentis* we identified 22,398 genes from 12,145 nuclei (548 genes per nucleus) and 23,979 genes from 15,786 protoplasts (1,945 genes per protoplast) (Fig. 1d-f). The Uniform Manifold Approximation and Projection (UMAP) algorithm was used to visualise local similarities and global structures of each dataset, and identified up to 19 clusters in the single-nucleus datasets and 15 clusters in the protoplast datasets (Fig. S4a-d). Transcriptome profiles were highly correlated between all replicates (Fig. S4e-h). Based on the expression of marker genes for leaf cell types we identified mesophyll, bundle sheath, xylem, phloem, epidermis, guard, and proliferating cell clusters in both single-nucleus and -protoplast datasets (Fig. 1 and Fig. S4i-l). We conclude that both methods captured sufficient numbers of cells, allowed identification of major cell types and the quantification of gene expression. As expected, fewer genes were detected in nuclei compared with protoplasts.

**Fig. 1.**
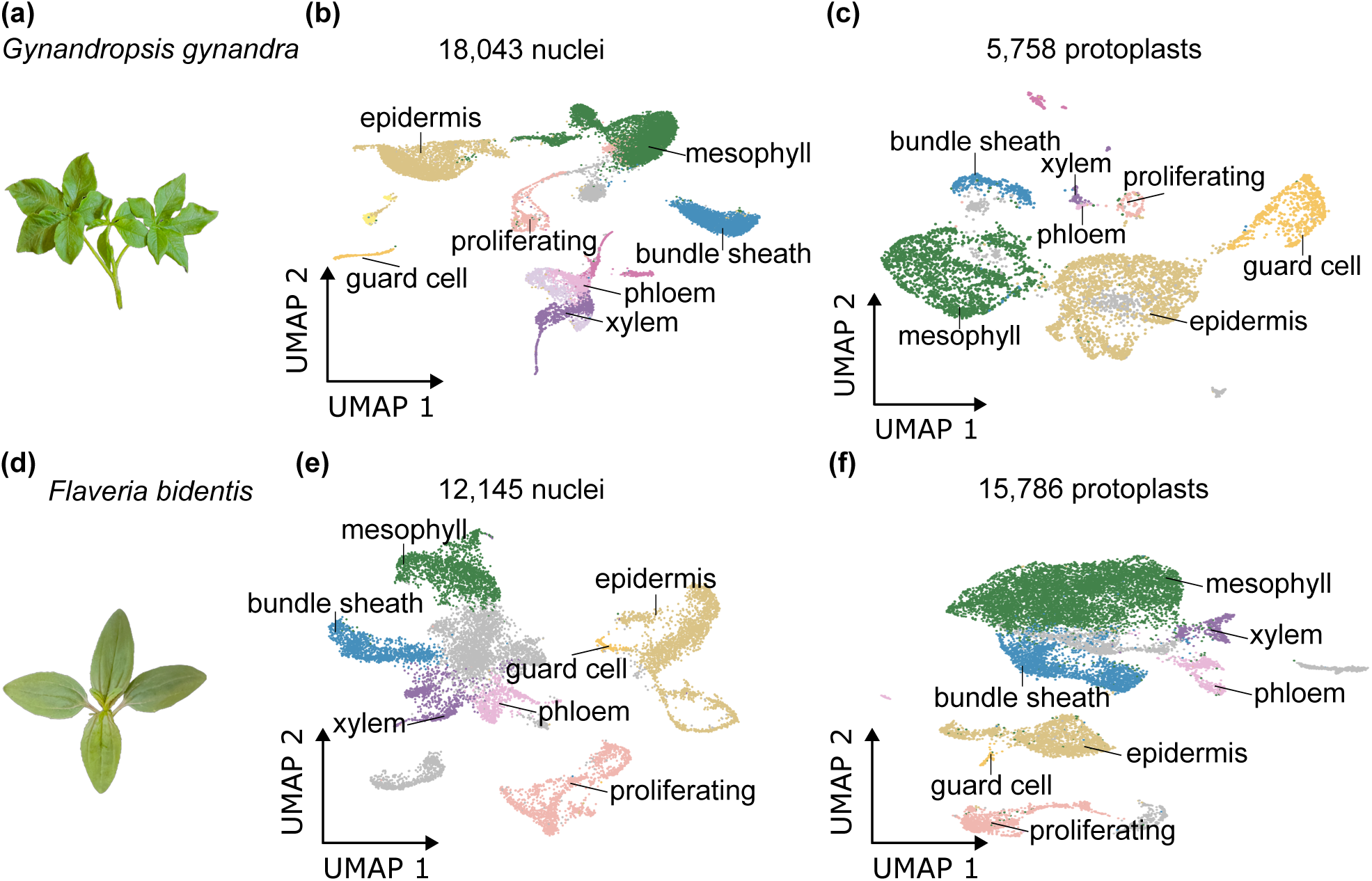
Single-nucleus and single-protoplast RNA sequencing from leaves of *Gynandropsis gynandra* and *Flaveria bidentis*. (a,d) Leaves of *G. gynandra* and *F. bidentis* were sampled for the isolation of single-nucleus and protoplasts for sequencing. Uniform Manifold Approximation and Projection (UMAP) visualisation of single-nucleus (b,e) and protoplast sequencing (c,f) from *G. gynandra* (b,c) and *F. bidentis* (d,e). Clusters coloured by cell identity.

### Leaf cell atlases for two distinct lineages of C_4_ plant

We integrated the single nucleus and protoplast datasets and re-clustered the combined dataset. This increased statistical power because more protoplasts/nuclei were available for each species. In *G. gynandra*, we detected the expression of 24,035 genes from 23,801 nuclei and protoplasts, and identified a total of 18 clusters. For *F. bidentis*, 24,874 genes were detected from 27,931 nuclei and protoplasts, and these could be separated into 21 clusters (Fig. S5a,b). Each cluster was annotated based on the differential expression of previously defined marker genes (Fig. 2a-d, Fig. S5c,d, and Table S3). In *G. gynandra* we identified all major leaf cell types, including mesophyll, bundle sheath, epidermis, guard cells, proliferating cells and vasculature cells (Fig. 2a,c). Mesophyll cells from *G. gynandra* (clusters 0, 5, and 8) showed elevated expression of such as *PHOSPHOENOLPYRUVATE CARBOXYLASE* (*PEPC*) and *PYRUVATE, ORTHOPHOSPHATE DIKINASE (PPDK*) that are needed for the C_4_ cycle. Bundle sheath cells (cluster 2) were identified based on strong expression of genes encoding the C_4_ acid decarboxylase NAD-dependent malic enzyme and transketolase from the Calvin Benson Bassham cycle. Expression of epidermis specific gene *KCS10* was predominantly detected in clusters 1, 4, 16, and 7. Cluster 7 not only exhibited high expression of epidermis marker genes but also showed enriched expression of guard cell marker genes such as *FAMA*. In previous studies, *HTA2* and *POK2* have been associated with cell proliferation during the cell cycle, especially during the synthesis phase and cytokinesis. In our dataset, we identified their expression mainly in clusters 10 and 15, indicating that these two clusters likely represent proliferating cells. Expression of the xylem-specific gene *ACAULIS5* (*ACL5*) was enriched in clusters 3 and 14, while cluster 3 also contained the phloem marker *TMO6*, suggesting that cluster 3 represents a mixed population of vascular cell types. Sub-setting and re-clustering cells from both cluster 3 and 14 allowed categorisation of phloem, xylem, and procambial cell types (Fig. S5a,c). We used similar marker genes (the only difference was the use of *NADP-ME* as a bundle sheath marker) to annotate cell types in *F. bidentis*, and this allowed us to annotate mesophyll (clusters 1, 2, 5, and 19), bundle sheath (clusters 3 and 8), xylem (cluster 9), phloem (cluster 15), epidermis (clusters 4, 7, 12, and 17), guard cells (cluster 18), and proliferating cells (clusters 6 and 14) (Fig. 2b,d, and Fig. S5b,d and D).

**Fig. 2.**
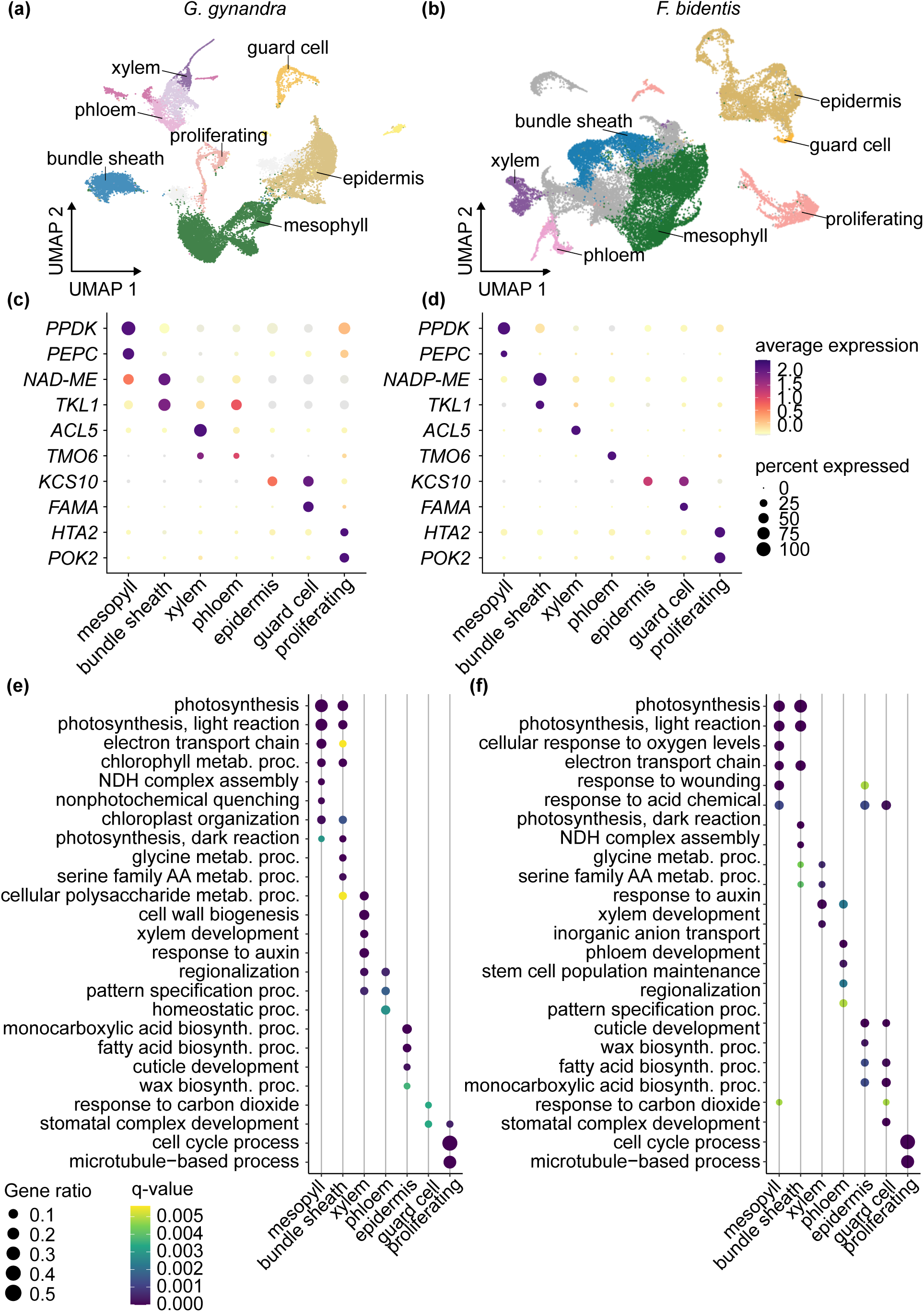
Integrated single-nucleus and single-protoplast atlases for *G. gynandra* and *F. bidentis* leaves. (a,b) UMAP visualization after integration of nucleus and protoplast data from *G. gynandra* (a) and *F. bidentis* (b). (c,d) Dot plots showing the expression of marker genes for major cell types of *G. gynandra* (c) and *F. bidentis* (d) leaves. Dot size represents the percentage of nuclei and protoplasts in each cluster expressing the gene indicated. Colour indicates the average expression level, calculated as the mean expression of the gene across all nuclei and protoplasts in which it is detected in that cluster. (e,f) Dot plots displaying enriched Gene Ontology (GO) terms derived from the marker genes of each annotated cell type cluster for *G. gynandra* (e) and *F. bidentis* (f). Dot size represents the fraction of marker genes associated with each GO term relative to the total number of input marker genes, and colour corresponds to q-value of enrichment. metb., short for metabolic; proc., short for process; biosynth., short for biosynthetic.

To gain insights into the functional specialisation of each cell type, we further analysed differentially expressed genes (DEGs) between all cell types (i.e. between one cell type and all others; p < 0.01; average log fold change ≥ 0.25; expressed in >10% of cells with a specific cell type) (Table S4, S5). DEGs with strong specificity (Fig. S6) included both the marker genes we had initially used to annotate cell types, but other genes that were also highly specific. For example, in *G. gynandra*, we identified 1,027 and 1,345 marker genes for the mesophyll and bundle sheath respectively, while in *F. bidentis* we found 2,209 and 1,257 for the mesophyll and bundle sheath. In both species, mesophyll markers included well-known genes associated with the C_4_ cycle such as *PPDK*, *PEPC* and photosystem subunits, while bundle sheath markers included genes required for the Calvin Benson Bassham cycle such as *RbcS* (*RIBULOSE BISPHOSPHATE CARBOXYLASE OXYGENASE SMALL SUBUNIT*), *FRUCTOSE-BISPHOSPHATE ALDOLASE1* (*FBA1*), or photorespiration including *GLD-T* (*GLYCINE DECARBOXYLASE T-SUBUNIT*), and *SHM1* (*SERINE HYDROXYMETHYLTRANSFERASE 1*) (Table S6, S7). We further performed gene ontology (GO) enrichment analysis using the top 200 DEGs of each cell type. In both species, the top 200 DEGs from both mesophyll and bundle sheath cells were significantly enriched in GO terms annotated as photosynthesis, light reactions, and electron transport, whilst those genes in bundle sheath cells were also enriched in terms associated with glycine metabolism and serine catabolism. Enrichment of marker genes for other cell types (xylem, phloem, epidermis, guard and proliferating cells) aligned with known function (Fig. 2e,f, Table S8, S9).

### Single-nucleus analysis captures more accurate cell-type composition and transcriptome profiles

Once data had been integrated, patterns of gene expression derived from nuclei and protoplasts as depicted by UMAPs showed reasonable congruence (Fig. 3a, Fig. S7a) and marker genes for each cell type exhibited similar patterns of expression (Fig. S7b, Fig. S8a). Thus, single-nucleus and single-protoplast sequencing captured comparable transcriptome profiles. However, whilst both approaches allowed us to detect all major cell types of the leaf including mesophyll, bundle sheath, epidermis, guard cells, proliferating cells, and vascular cells, the abundance of these cell types differed between the two approaches. For example, although a similar percentage of cells detected were annotated as mesophyll, nuclei had a greater representation of internal cells such as vascular cells and the bundle sheath, and much lower proportions of guard and epidermal pavement cells compared with protoplasts (Fig. 3b). Notably, in *G. gynandra* the most abundant cell type detected after protoplasting was the epidermis (43% of all cells). Similar trends were observed for *F. bidentis* (Fig. S7c). Although some differences in cell type composition between datasets may be explained by species differences or different developmental stage, in general the single-nucleus datasets appeared to more accurately reflect the cell-type composition of a leaf and captured internal cells that are less likely to be digested by cell wall degrading enzymes.

**Fig. 3.**
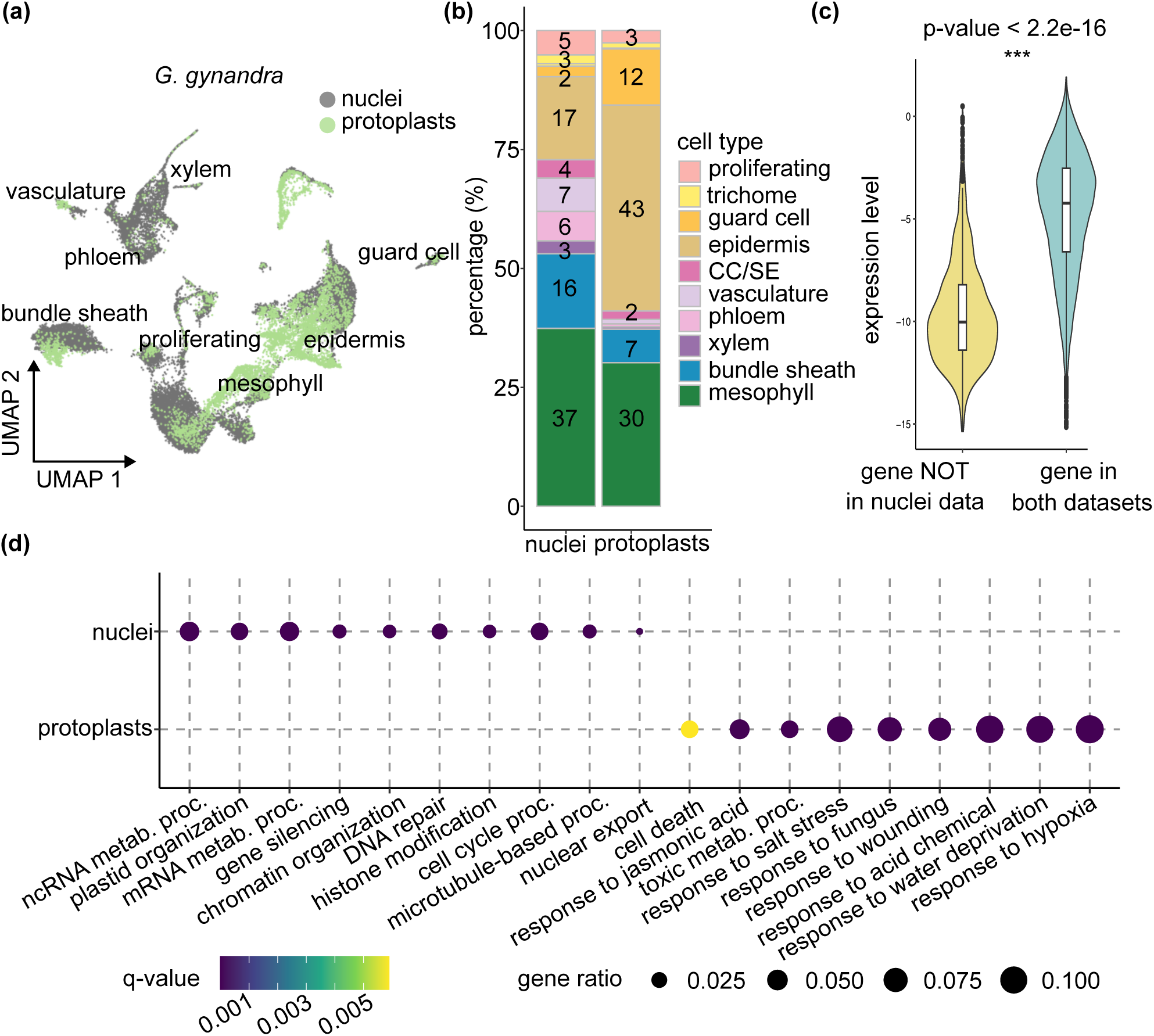
Comparison of single-nucleus and single-protoplast transcriptomes from *G. gynandra* leaves. (a) UMAP visualization of combined protoplast and nucleus transcriptomes. Major clusters annotated by cell-type. Nuclei shown as grey and protoplasts as green. (b) Bar chart showing representation of each cell type estimated from nucleus or protoplast data based on expression of marker genes. CC/SE, companion cell/sieve element related cell. (c) Violin plot illustrating expression of genes detected only in protoplasts (yellow) or both nuclei and protoplasts (teal). (d) Dot plots showing enriched Gene Ontology (GO) terms for genes that were differentially expressed between nuclei and protoplasts. metb., short for metabolic; proc., short for process.

Fewer genes were detected in nuclei compared with protoplasts, but this effect could be offset by sequencing more nuclei such that the total number of genes detected was similar between these datasets (Fig. S7d, Fig. S8b). Genes not detected by single-nucleus sequencing were primarily poorly expressed genes (Fig. 3c, Fig. S7e,f, Fig. S8c). Analysis of differences in transcript abundance between nuclei and protoplasts in *G. gynandra* highlighted that the nuclei data mostly captured genes associated with RNA, DNA organisation and gene expression, whilst protoplast data captured many genes related to stress responses (Fig. 3d, Fig. S9a,c and C, Table S10, S11. This was also the case in *F. bidentis* (Fig. S7h, Fig. S8b,d, Table S12, S13). Overall, we conclude that both single nucleus and protoplast data allow identification of major cell types from C_4_ leaves, but analysis of nuclei appeared to provide two advantages. First, the representation of different cell types was more comparable to the proportion of each cell type found in the leaf, and second there was less evidence for upregulation of stress response pathways. We therefore chose to analyse the single-nucleus data from *G. gynandra* and *F. bidentis* to better understand similarities and differences in gene expression between these C_4_ plants.

### Conserved patterning of structural genes as well as transcription factors in two different C₄ subtypes

To provide context for analysis of the single nucleus data from *G. gynandra* and *F. bidentis* we first compared each with publicly available data from the C_3_ model *Arabidopsis thaliana* (Kim et al., 2021). In both cases, comparison of marker genes for each cell type indicated a significant amount of conservation between the patterns of gene expression in cell type (Fig. 4a,b). However, bundle sheath cells from *G. gynandra* and *F. bidentis* shared greater similarity with mesophyll cells from *A. thaliana* than with the bundle sheath (Fig. 4c), and were more similar to each other than to these cells from C_3_ *A. thaliana* (Fig. 4a-c). This similarity was associated with genes involved in the light-dependent reactions of photosynthesis (e.g. LHCA1: OG0009556, LHCA3: OG0009361, and LHCA4: OG0007442), the Calvin Benson Bassham cycle (e.g. FBA: OG0001032), and photorespiration (including subunits of the glycine decarboxylase complex, GLDT: OG0008599 and GLDH: OG0008404, as well as SHM1: OG0002906) - see Supplementary table 4 and table 5 for cell type markers, and Supplementary table 14 for their orthology. Thus, despite diverging from a common ancestor around 100 million years ago, and employing different C_4_ acid decarboxylases to release CO_2_ around RuBisCO, *G. gynandra* and *F. bidentis* exhibited similar patterns of gene expression in bundle sheath cells.

**Fig. 4.**
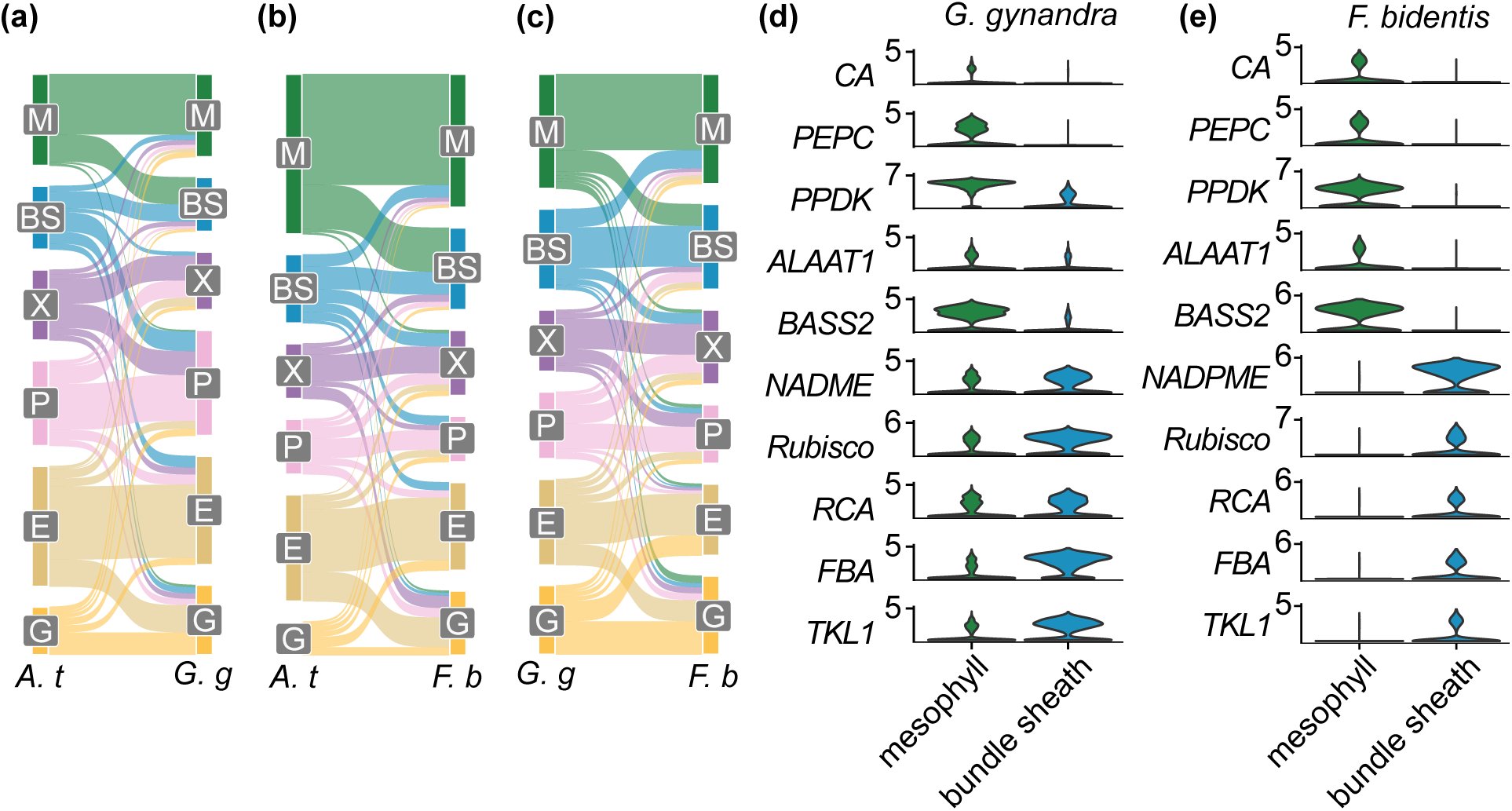
Compartmentalization of genes between bundle sheath and mesophyll. (a-c) Sankey plots summarising changes in compartmentation of gene expression between C_3_ *Arabidopsis thaliana* and C_4_ *G. gynandra* (a), C_3_ *A. thaliana* and C_4_ *F. bidentis* (b), and C_4_ *G. gynandra* and C_4_ *F. bidentis* (c). *A. t.*, *Arabidopsis thaliana*; *G. g.*, *Gynandropsis gynandra; F. b*., *Flaveria bidenti*s. (d,e) Violin plots showing transcript abundance of C_4_ cycle and Calvin-Benson-Bassham (CBB) cycle genes in mesophyll (green) and bundle sheath (blue) cells of *G. gynandra* (d) and *F. bidentis* (e).

As would be expected, core C_4_ genes showed similar patterns of expression in both species (Fig. 4d,e), and photorespiration genes were rewired to be strongly expressed in the bundle sheath (Fig. S10). Whereas expression of genes encoding enzymes of photorespiration need to be downregulated in C_4_ plants compared with the ancestral C_3_ state, those of the C_4_ cycle are upregulated in either mesophyll or bundle sheath cells. Most C_4_ cycle genes are encoded by multigene families and so we investigated whether particular gene copies are recruited into C_4_ photosynthesis in both *G. gynandra* and *F. bidentis*. For most proteins, in both species gene copies from one orthogroup were upregulated in either mesophyll or bundle sheath (XXX). This included genes encoding a chloroplastic PYROPHOSPHATASE from OG0007017 needed for PPDK to function in mesophyll cells (Fig. S11,12 and Table S15). However, we also found evidence that convergent evolution underpins recruitment of C_4_ cycle enzymes in these species. For example, for both *PHOSPHO*ENOL*PYRUVATE CARBOXYLASE (PEPC)* and *ADENYLATE KINASE (AMK)* different gene copies became strongly expressed in mesophyll cells of *G. gynandra* and *F. bidentis* (Fig. S13,14). Moreover, in the case of *CARBONIC ANHYDRASE (CA)* where previous analysis indicated loss of a chloroplast transit peptide allowed increased activity in the cytosol of mesophyll cells of *Flaveria* (Tanz, Tetu, Vella, & Ludwig, 2009) and also *Neurachne* (Clayton et al 2017), in *G. gynandra* we found no evidence for this (Fig. S15 and Table S16). Instead, in *G. gynandra* transcripts encoding a cytosolic CA isoform were upregulated in mesophyll cells.

The data also allowed bundle sheath and other vascular cells to be separated from each other, and so provide insight into patterns of gene expression specifically associated with the bundle sheath. For example, in both species, genes preferentially expressed in the mesophyll were also generally highly expressed in epidermis and guard cells, whereas genes preferentially expressed in the bundle sheath were often highly expressed in the xylem or phloem (Fig S16). Previous work established that the three C_4_ acid decarboxylases (NADP-ME, NAD-ME and PEPCK) that provide CO_2_ to RuBisCO in C_4_ plants are preferentially expressed in vascular cells of C_3_ species (Hibberd and Quick, 2002; Brown et al 2010). Here we investigated whether expression of these genes was detected in phloem and/or xylem cells of *G. gynandra* and *F. bidentis*. In *G. gynandra*, both *NAD-ME* genes showed strong expression in the bundle sheath, and *NAD-ME1* was also detected in phloem cells. In *F. bidentis* one *NADP-ME* gene was strongly expressed in the bundle sheath, but was barely detected in the vasculature (Fig. S17 and Fig. S18). *PEPCK* did not show strong expression in *G. gynandra* bundle sheath, but was detected in the phloem (Fig. S19).

To better understand how signatures of bundle sheath and mesophyll cells may be specified we next explored transcription factors that were preferentially expressed in each cell type. 38 mesophyll and 44 bundle sheath preferential transcription factors were identified in *G. gynandra*, whilst in *F. bidentis* 32 mesophyll and 37 bundle sheath preferential transcription factors were detected (Fig. 5a). As with marker genes, transcription factors preferentially expressed in the mesophyll were also highly expressed in the epidermis and guard cells, while those enriched in the bundle sheath exhibited stronger expression in vascular tissues. Families such as the bHLH, C2H2, and homeobox transcription factors had genes that were specific to mesophyll and bundle sheath cells in both *G. gynandra* and *F. bidentis* (Fig. 5b). However, it was notable that in both species members of the DOF and IDD families were specific to bundle sheath cells (Fig. 5b and Fig. S20). These were also identified as bundle sheath specific from the protoplast data (Fig. S21). Moreover, we identified fifteen transcription factors belonging to the same orthogroups that were specific to the either mesophyll or bundle sheath cells from both species (Fig. 5c). This included seven that were bundle sheath preferential and eight that were mesophyll preferential. The top two most strongly expressed of these in the bundle sheath (BRON and IDD9) belonged to the C2H2 family.

**Fig. 5.**
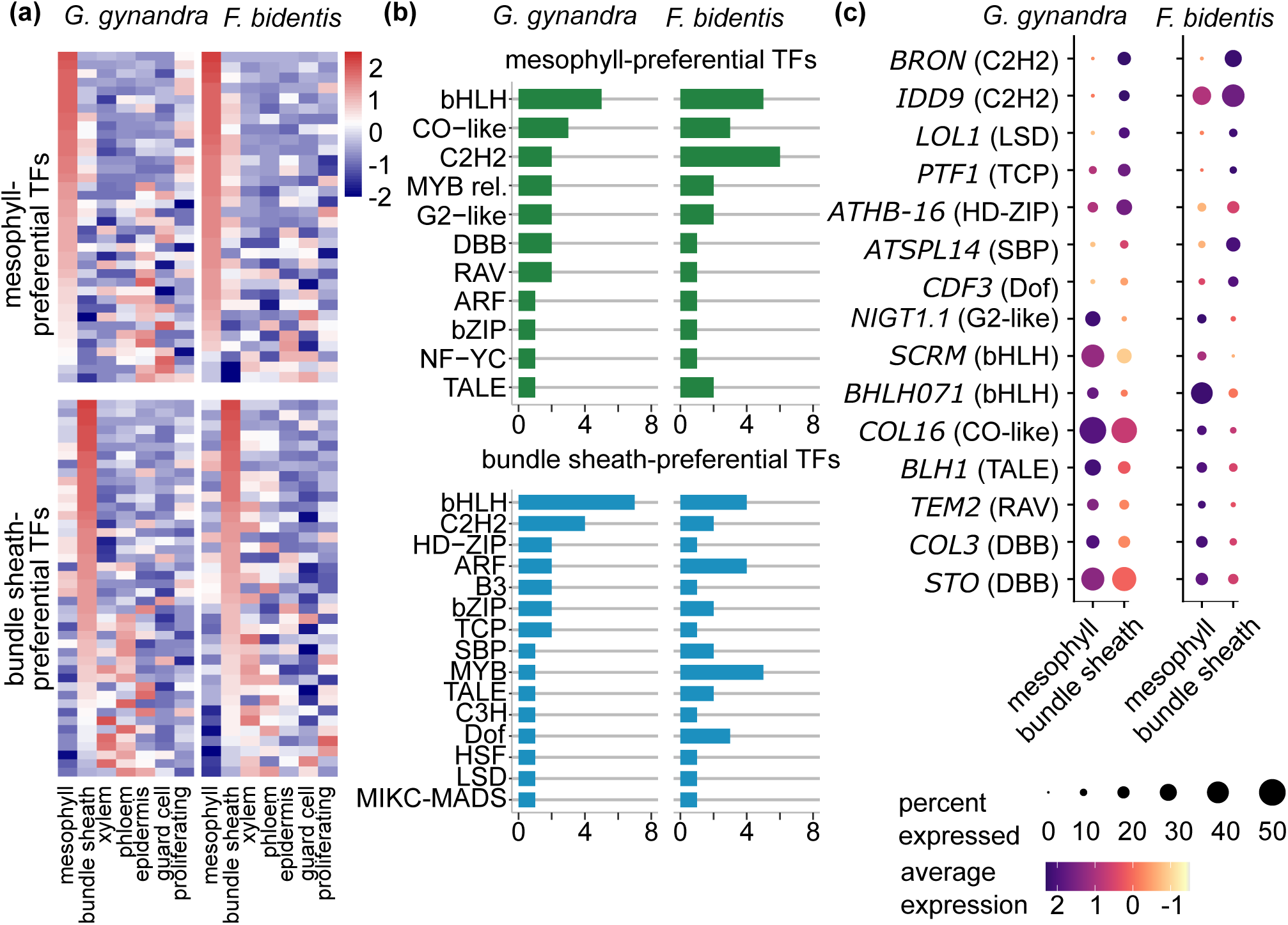
Conserved patterning of transcription factors between bundle sheath and mesophyll cells. (a) Heatmap of transcription factors that were differentially expressed between mesophyll and bundle sheath cells of *G. gynandra* (left) and *F. bidentis* (right). (b) Number of transcription factors from families that were preferentially expressed in either mesophyll or bundle sheath cells of both *G. gynandra* (left) and *F. bidentis* (right). (c) Transcript abundance of orthologous transcription factors that were preferentially expressed in either mesophyll or bundle sheath cells of both *G. gynandra* (left) and *F. bidentis* (right).

In summary, we report optimization of both single-nucleus and single-protoplast RNA-SEQ and as a consequence construct high-resolution transcriptome atlases for two independently evolved C_4_ species. Our analysis indicates that nucleus sequencing better preserves native cell-type composition, and transcriptome profiles showed lower signatures of stress. Using the nuclei data, we confirmed core C_4_ gene compartmentalization and identified transcriptional signatures of mesophyll and bundle sheath cells. Further we found that in both species, mesophyll-enriched genes were often co-expressed in epidermal and guard cells, suggesting some degree of shared regulation between these cell types. In contrast, bundle sheath-enriched genes overlapped more with xylem and phloem, pointing to a closer transcriptional relationship with these vascular tissues. Together, our findings provide new insights into the spatial partitioning of gene expression needed for C_4_ photosynthesis and serve as a valuable resource for future studies on the evolution and engineering of C_4_ traits.

## Discussion

After direct comparison of single protoplast and single nucleus sequencing, we propose that the latter provides a more accurate representation of cell type composition and more effectively characterizes native gene expression across different cell types from leaves. Both approaches validated known patterns of gene expression associated with C_4_ photosynthesis. However, by capturing transcriptomes at single-cell or single-nucleus resolution, we were also able to separate signatures from bundle sheath cells and vascular tissues, and by including two independently evolved C_4_ species identified transcription factors not previously associated with bundle sheath cells. These cell specific atlases provide valuable resources to study gene compartmentalization in the context of leaves in general and C_3_ and C_4_ photosynthesis in particular.

To generate high-quality transcriptome atlases at cellular resolution for leaves of two C_4_ dicotyledons, we employed various methods to optimize protoplast and nuclei isolation protocols tailored for the most widely used single-cell platforms. As previously reported (Chupeau et al., 2013; Sawers et al., 2007) protoplast isolation led to transcriptome signatures associated with stress. Because of this sensitivity of leaf transcriptomes to abiotic stress during protoplast isolation, despite nuclei-based analysis detecting fewer transcripts, we conclude that their analysis is informative as stress-related signals are lower, and that the proportion of each cell type detected is closer to tissue composition of the leaf. Because of these advantages we primarily focussed on analysis of the nuclei data to investigate gene expression in the two C_4_ dicotyledons *G. gynandra* and *F. bidentis*. In contrast to C_4_ species in the monocotyledons (Bezrutczyk et al., 2021; Swift et al., 2024), although both *G. gynandra* and *F. bidentis* are used as models to understand C_4_ photosynthesis (Brown et al., 2011; Gowik et al., 2011; Hoang et al., 2023; Schreier et al., 2024; Schulze et al., 2013; Singh et al., 2023) single-cell data have previously not been available. Thus, our analysis provides transcriptome data not only for bundle sheath and mesophyll cells but also for other critical cell types, such as vascular and guard cells, which play essential roles in the C_3_ and C_4_ leaves. Although these two C_4_ species diverged from a common ancestor over 100 million years ago (Moore et al., 2010; Zuntini et al., 2024), we find that the transcriptome profile of bundle sheath cells in *G. gynandra* shares greater similarity with *F. bidentis* than with those available from the more closely related C_3_ model, *A. thaliana*, which diverged from a common ancestor with *G. gynandra* 41 million years ago (Schranz & Mitchell-Olds, 2006). Previous single nucleus analysis of the monocotyledons rice and sorghum revealed that all cell types showed differences in gene expression between these C_3_ and C_4_ species, but the bundle sheath cells showed the most significant rewiring (Swift et al., 2024). Thus, combined with this previous work, the data reported here indicate that evolution of the C_4_ pathway in the dicotyledons as well as the monocotyledons has resulted in major rewiring of the bundle sheath. Even when distinct biochemistries are being used to decarboxylate CO_2_ in the bundle sheath cells of these two species, their transcriptomes showed greater convergence to each other than to bundle sheath cells from C_3_ *A. thaliana*.

To better understand the basis for this convergence in bundle sheath gene expression in *G. gynandra* and *F. bidentis*, we investigated transcription factors that were preferentially expressed in these cells in both species. To our knowledge there are only two examples where transcription factors that are both necessary and sufficient for bundle sheath expression have been defined. The first comprises a bipartite MYC-MYB complex in *A. thaliana* (P. J. Dickinson et al., 2020) that has now been implicated in the evolution of C_2_ photosynthesis (Patrick J. Dickinson, Triesch, Schlüter, Weber, & Hibberd, 2023). The second includes a collective of transcription factors from WRKY, G2-like, DOF, MYB-related, IDD and bZIP families in rice (Hua et al., 2024; Swift et al., 2024). Interestingly, in the data we report here members of transcription factors from the MYB but not the MYC family were bundle sheath preferential in both *F. bidentis* and *G. gynandra.* This is the same patterning of expression reported for *A. thaliana* where MYBs rather than MYCs drive the cell specificity (P. J. Dickinson et al., 2020). Moreover, although first reported in monocotyledonous species (Hua et al., 2024; Swift et al., 2024) the single nucleus sequencing reported here now reveals that transcription factors from the DOF, IDD and bZIP families are preferentially expressed in bundle sheath cells of *F. bidentis* and *G. gynandra*. In rice, DOF and IDD motif also enriched in the bundle sheath (Swift et al., 2024), thus, it is possible that a team of transcription factors based on these core members mediates bundle sheath expression in dicotyledons. In addition to these previously reported regulators of bundle sheath expression, we also identified previously unknown transcription factor families and orthogroups that were preferentially expressed in this cell type in both *F. bidentis* and *G. gynandra.* This included C2H2, LSD, HD-ZIP, and SBP transcription factors. Currently available single nucleus data from C_3_ models in the dicotyledons such as *A. thaliana* (Kim et al., 2021; Lopez-Anido et al., 2021; Tenorio Berrio et al., 2022) have not been performed at sufficient depth to determine if these genes are also preferentially expressed in bundle sheath cells in the ancestral C_3_ state. Furthermore, although transformation of *F. bidentis* and *G. gynandra* is possible it is relatively laborious (Chitty, Furbank, Marshall, Chen, & Taylor, 1994; Newell et al., 2010), and so functional analysis confirming an unequivocal role of these transcription factors in cell specificity will need to be proved in the future. To address these questions future studies that obtain single nucleus RNA-seq from close C_3_ relatives of these species will be helpful. Moreover, analysis of, for example the *Flaveria* genus, that contains intermediate photosynthetic types (that appear to span a trajectory from C_3_, to C_3_-C_4_, to C_2_ and C_4_ photosynthesis) (R. F. Sage, Sage, & Kocacinar, 2012) are also likely to inform us about rewiring of both mesophyll and bundle sheath cells during the evolution of C_4_ photosynthesis. Recent work using single-nucleus RNA-seq of a C_3_–C_4_ intermediate, *Moricandia arvensis,* highlighted functional specialization of bundle sheath cells in the photorespiratory shuttle, which exemplifies the power of such approaches (Triesch et al., 2024).

In summary, our single cell resolution transcriptome atlases for *G. gynandra* and *F. bidentis* leaves provide the first comprehensive maps for gene expression at this level for C_4_ dicotyledons. They capture all major cell types in the leaf, and also provide the quantification of gene expression in each of these cell types. To maximize the accessibility and utility of this atlas, we developed a web-based portal that is user-friendly and interactive for data visualization and exploration. The portal includes side-by-side UMAP plots to allow comparison of gene expression and cell annotation, proportion plots to assess cell group composition, and dot plots and heatmaps to visualise transcripts derived from multiple genes across selected cell groups. By facilitating efficient exploration of these complex datasets, our web portal empowers the research community to fully utilize and build upon the atlas.

## Acknowledgements

The work was funded by European Union Program (project GAIN4CROPS GA number 862087) to JMH and APMW and European Research Council Advanced Grant 694733 REVOLUTION to JMH. For the purpose of open access, the authors have applied a Creative Commons Attribution (CC BY) license to any Author Accepted Manuscript version arising from this submission. The authors thank Joseph Swift for helpful discussions regarding the single-nucleus sequencing experiments and analyses.

## Author contributions

JMH, LL, and TS conceived and directed the research. TS, LL, and JMH designed the experiments. TS and LL performed the single-protoplast and single-nucleus RNA-seq experiments. TS analyzed the single-protoplast and single-nucleus RNA-seq data. VSB and BS performed the genome-related experiments and analyses. TS, LL, and JMH wrote the manuscript with input from all authors.

## Data and code availability

All data generated in this study are publicly available. The genome sequence data reported in this paper are available at NCBI under BioProject ID PRJNA1259992, and the genome assembly and annotation can be accessed at: https://figshare.com/s/7fb9444d7cefef0c341f. The single-protoplast and single-nucleus sequencing data are available at ArrayExpress under accession number E-MTAB-15121. Our atlases can be explored via online web portals: https://tianshu-sun.shinyapps.io/gg_integrated/ for *G. gynandra* and https://tianshu-sun.shinyapps.io/fb_integrated/ for *F. bidentis*. The code generated during this study is available at GitHub: https://github.com/TianshuSunCam/C4_dicots_single_nucleus_protoplast. Further details needed to reanalyze the data in this study are available from the lead contact upon reasonable request.

## Supporting information

### Supplementary Figure

**Fig. S1** Workflow illustrating sampling and isolation of nuclei (left) and protoplasts (right) for sequencing.

**Fig. S2** Representative FACS plots of nuclei sorting.

**Fig. S3** Optimization of protoplast isolation for single-protoplast RNA-seq.

**Fig. S4** Clustering and cell type annotation of single-nucleus and single-protoplast RNA sequencing from leaves of *G. gynandra* and *F. bidentis*.

**Fig. S5** Cluster annotation for the integrated transcriptome atlases of *G. gynandra* and *F. bidentis* leaves.

**Fig. S6** Number of cell type marker genes shared across different cell types.

**Fig. S7** Comparison of single-nucleus and single-protoplast transcriptome profiles in *F. bidentis* leaves.

**Fig. S8** Comparison of gene detection between single-nucleus and single-protoplast transcriptome profiles from *G. gynandra* leaves.

**Fig. S9** Comparison of stress and nuclei signature scores.

**Fig. S10** Expression of photorespiration genes.

**Fig. S11** Phylogenetic reconstruction of the species used to identify orthogroups.

**Fig. S12** Phylogenetic tree of the PPA orthogroups.

**Fig. S13** Phylogenetic tree of the PEPC orthogroup.

**Fig. S14** Phylogenetic tree of the AMPK orthogroup.

**Fig. S15** Phylogenetic tree of the CA orthogroup.

**Fig. S16** Expression of differentially expressed genes between mesophyll and bundle sheath.

**Fig. S17** Phylogenetic tree of the NAD-ME orthogroups.

**Fig. S18** Phylogenetic tree of the NADP-ME orthogroup.

**Fig. S19** Phylogenetic tree of the PEPCK orthogroup.

**Fig. S20** Number of all mesophyll and bundle sheath preferential transcription factors characterised into different families in *G. gynandra* and *F. bidentis*.

**Fig. S21** Compartmentation of transcription factors between bundle sheath and mesophyll using protoplast data only.

### Supplementary Tables

Table S1. Assessment of contiguity and completeness of the reference-quality assembly

Table S2. Amount of genome sequence data generated using each type of sequencing platform

Table S3. List of marker genes used for cell type annotation

Table S4. List of marker genes for each cell type in *G. gynandra*

Table S5. List of marker genes for each cell type in *F. bidentis*

Table S6. Best BLAST hit of *G. gynandra* genes in *A. thaliana*

Table S7. Best BLAST hit of *F. bidentis* genes in *A. thaliana*

Table S8. GO enrichment analysis of *A. thaliana* orthologs of *G. gynandra* cell type marker genes.

Table S9. GO enrichment analysis of *A. thaliana* orthologs of *F. bidentis* cell type marker genes.

Table S10. List of differentially expressed genes between protoplast (avg_log2FC>0) and nuclei in *G. gynandra*

Table S11. GO enrichment analysis of *A. thaliana* orthologs of differentially expressed genes between protoplasts and nuclei in *G. gynandra*

Table S12. List of differentially expressed genes between protoplast (avg_log2FC>0) and nuclei in *F. bidentis*

Table S13. GO enrichment analysis of *A. thaliana* orthologs of differentially expressed genes between protoplasts and nuclei in *F. bidentis*

Table S14. Orthogroup description for top marker genes shared between *G. gynandra* and *F. bidentis* (deduplicated)

Table S15. TargetP prediction of the subcellular localization of PPAs Table S16. TargetP prediction of the subcellular localization of CAs

