## Supporting information for "Conserved spatial patterning of gene expression in independent lineages of C_4_ plants"

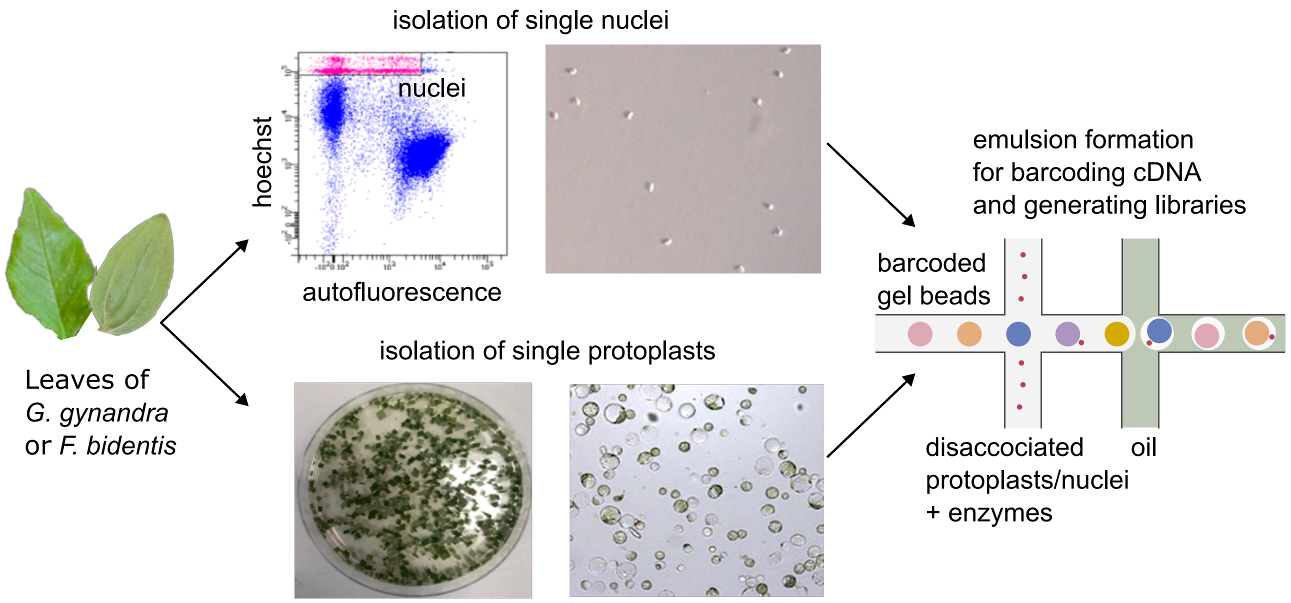


**Fig. S1**

Workflow illustrating sampling and isolation of nuclei (left) and protoplasts (right) for sequencing. Leaves from *G. gynandra* and *F. bidentis* were collected, prior to nuclei isolation through Fluorescence-activated cell sorting (FACS, top) or protoplast isolation through enzymatic digestion (bottom). Isolated nuclei and protoplasts were then subjected to single-cell RNA sequencing for downstream analysis.

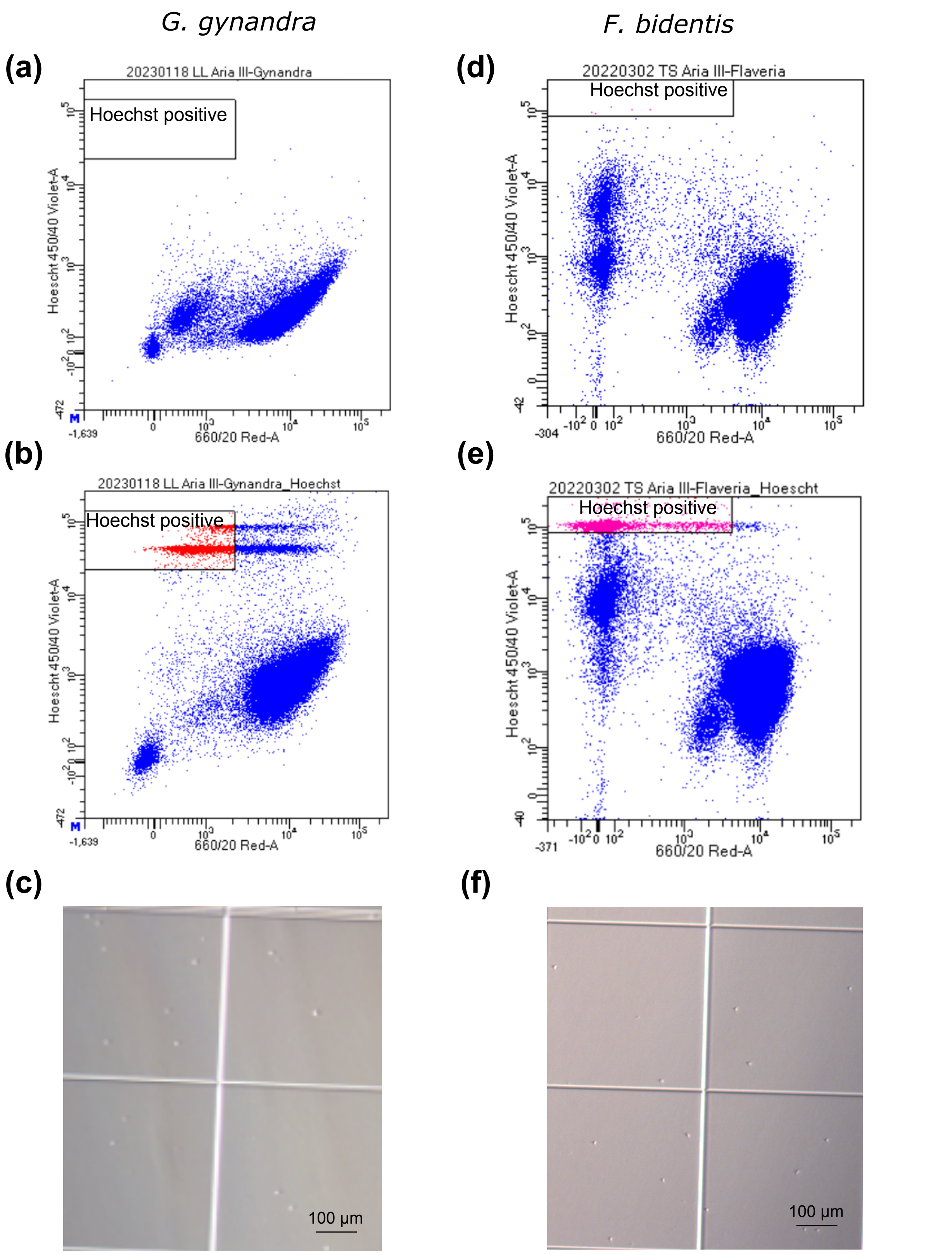


**Fig. S2**

Representative FACS plots of nuclei sorting. The gate for unstained *G. gynandra* nuclei (a) and *F. bidentis* nuclei (d) showing Hoechst fluorescence signals on the y-axis plotted against autofluorescence signals on the x-axis. The gate for Hoechst+ *G. gynandra* nuclei is shown in red (b) and for Hoechst+ *F. bidentis* nuclei in magenta (e). (c) and (f) Images of sorted nuclei under a light microscope.

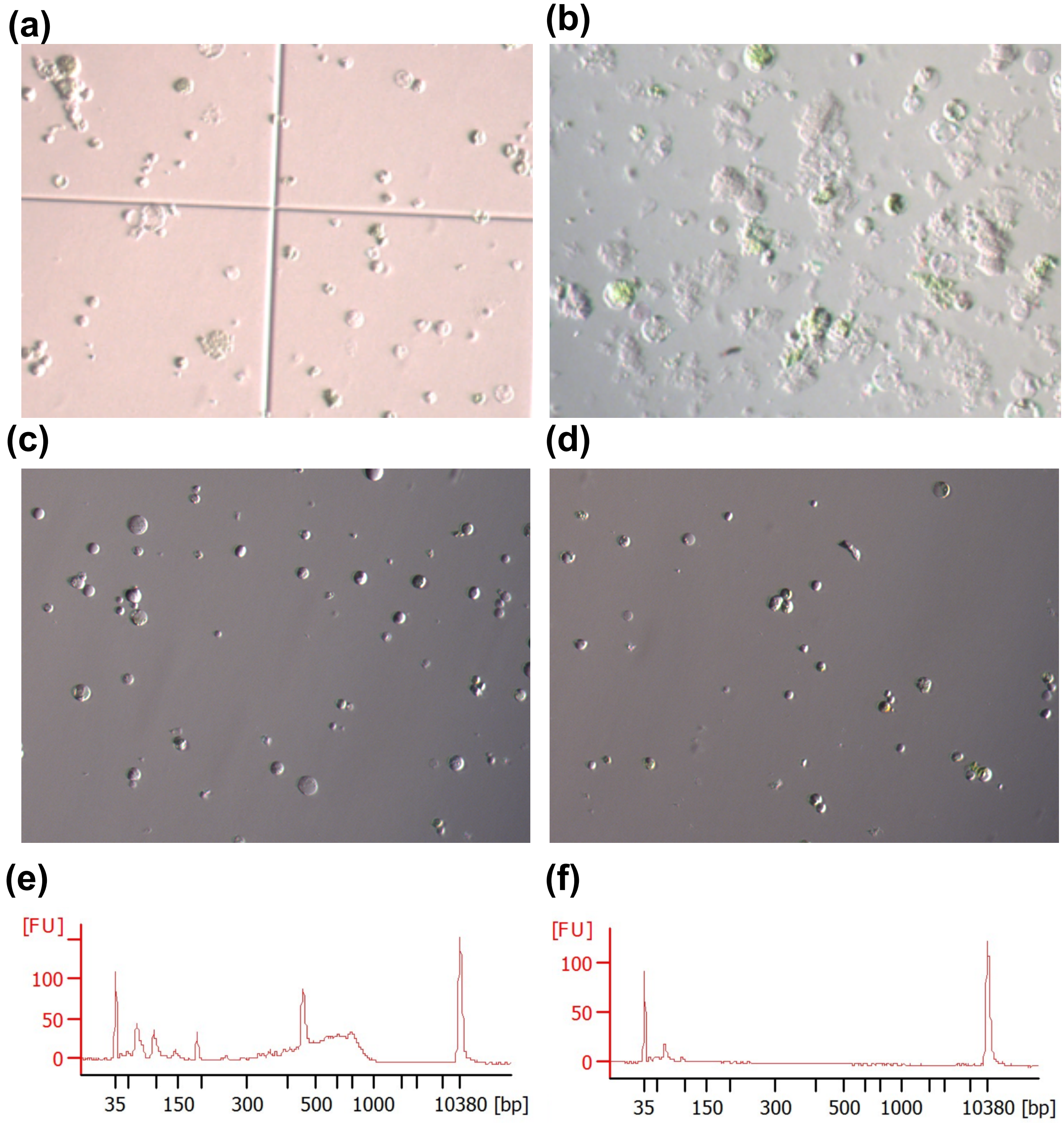


**Fig. S3**

Optimization of protoplast isolation for single-protoplast RNA-seq. Light microscopy images show isolated *F. bidentis* protoplasts derived from young leaves of a 5-week-old plant (a) and a 2-week-old plant (b). Isolated protoplasts resuspended in mannitol (c) or ½ W5 buffer (d) shown. Successful cDNA synthesis and amplification were achieved in mannitol-resuspended protoplast samples (e), whereas no cDNA was amplified in W5-resuspended samples.

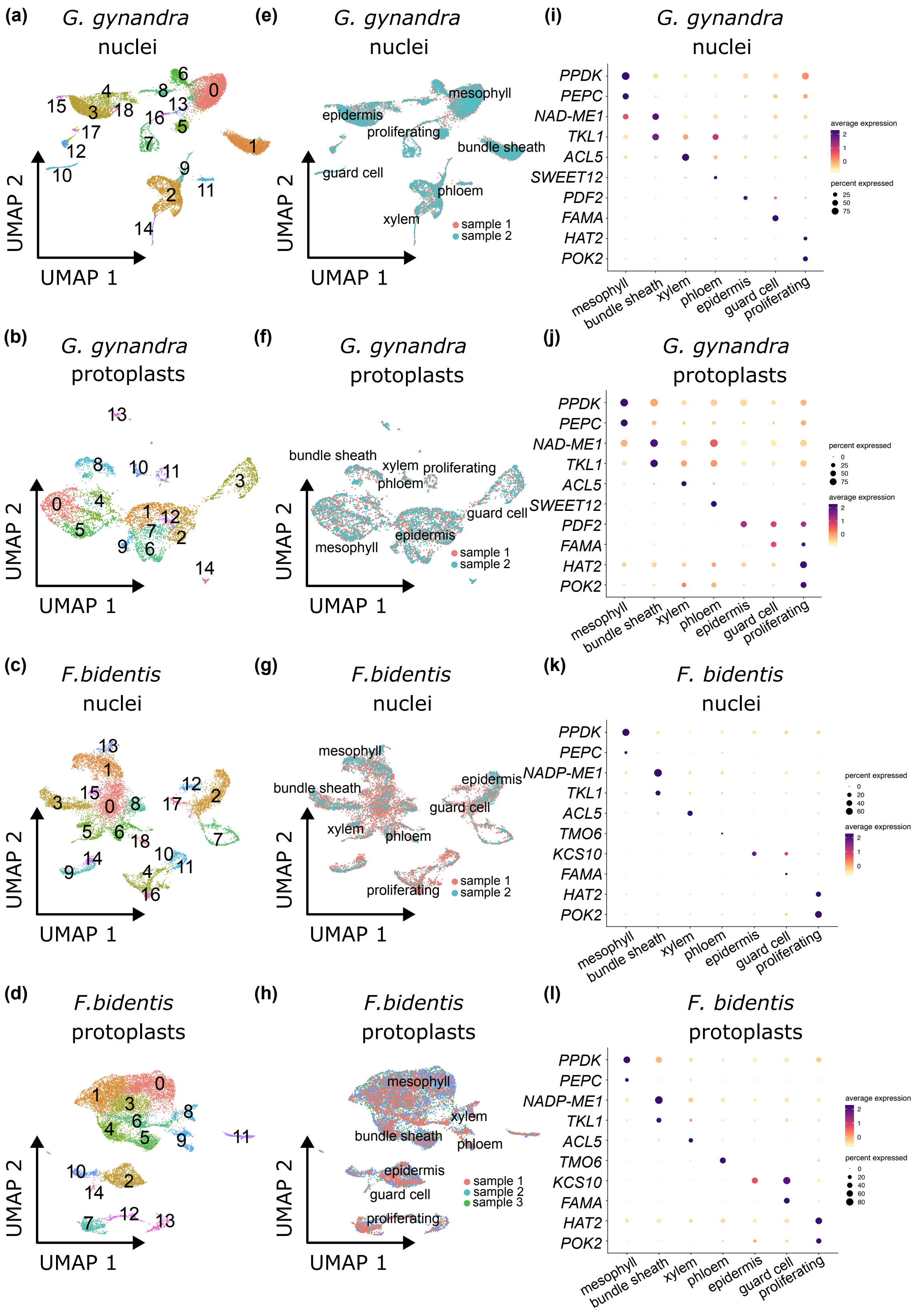


### Fig. S4

Clustering and cell type annotation of single-nucleus and single-protoplast RNA sequencing from leaves of *G. gynandra* and *F. bidentis*. (a-d) UMAP visualisation of the clustering of nuclei or protoplasts from *G. gynandra* (a) and (b) or *F. bidentis* (c) and (d), coloured by unsupervised clusters. (e-h) UMAP visualisation of the clustering of nuclei or protoplasts from *G. gynandra* (e) and (f) or *F. bidentis* (g) and (h), coloured by different replicates. (i-l) Dot plots showing expression marker genes defining major annotated cell types.

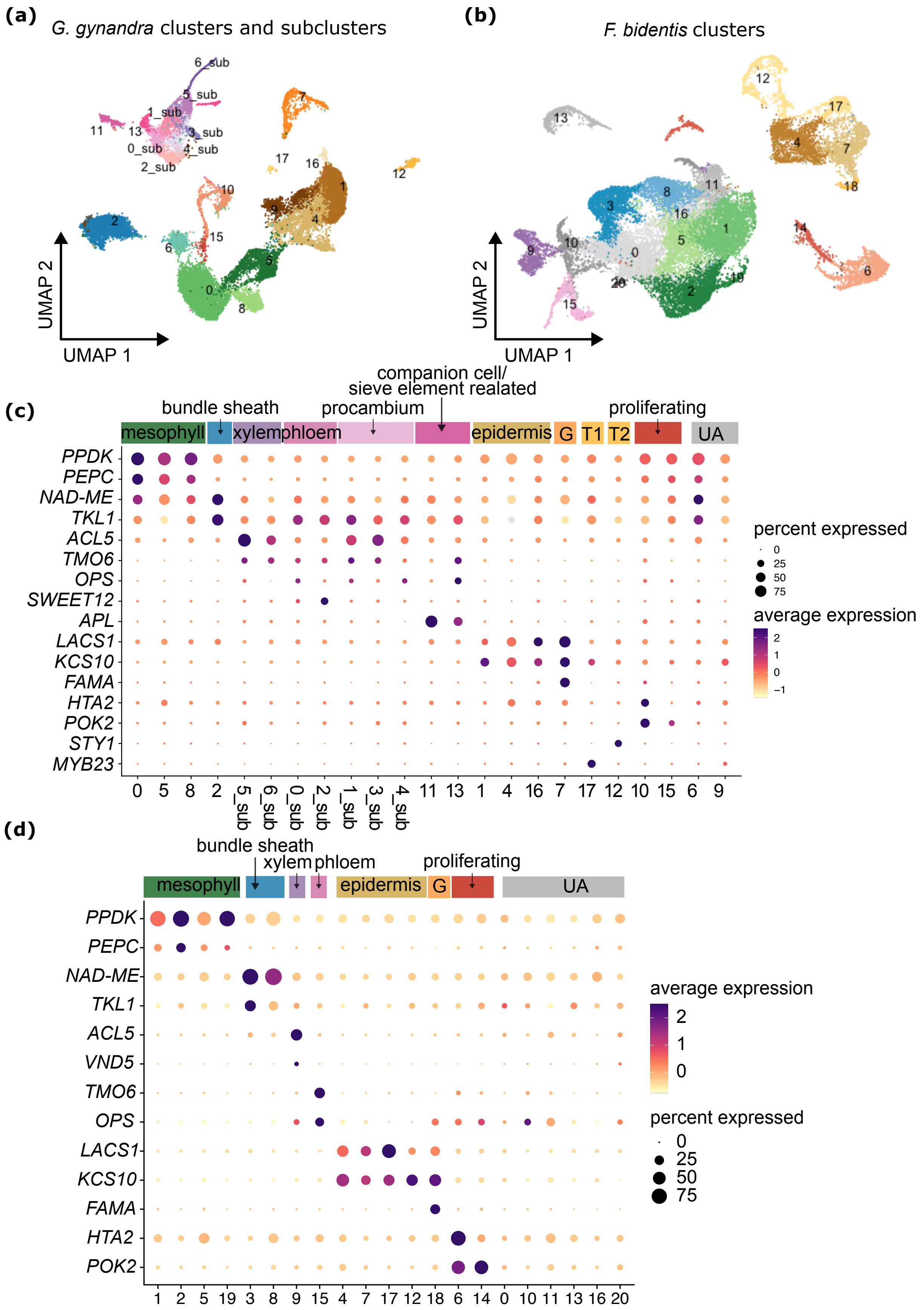


### Fig. S5

Cluster annotation for the integrated transcriptome atlases of *G. gynandra* and *F. bidentis* leaves. UMAP visualization of combined nuclei and protoplast transcriptome profiles of *G. gynandra* (a) and *F. bidents* (b), coloured by unsupervised clusters. Dot plots showing the expression of cell type-defining marker genes for each cluster in *G. gynandra* (c) and *F. bidents* (d).

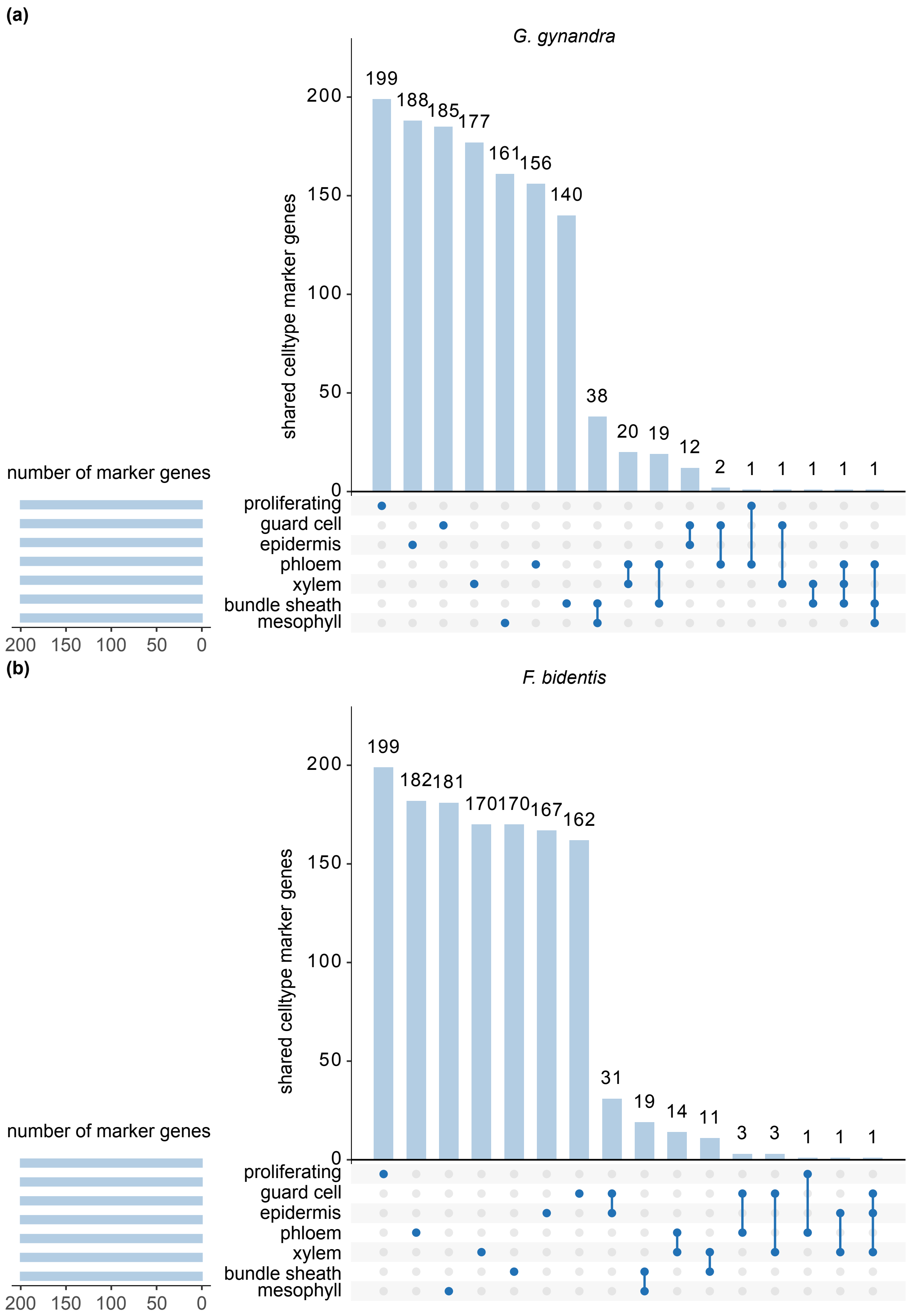


**Fig. S6**

Number of cell type marker genes shared across different cell types. Upset plots illustrating the unique and shared top 200 marker genes across different cell types in *G. gynandra* (a) and *F. bidentis* (b).

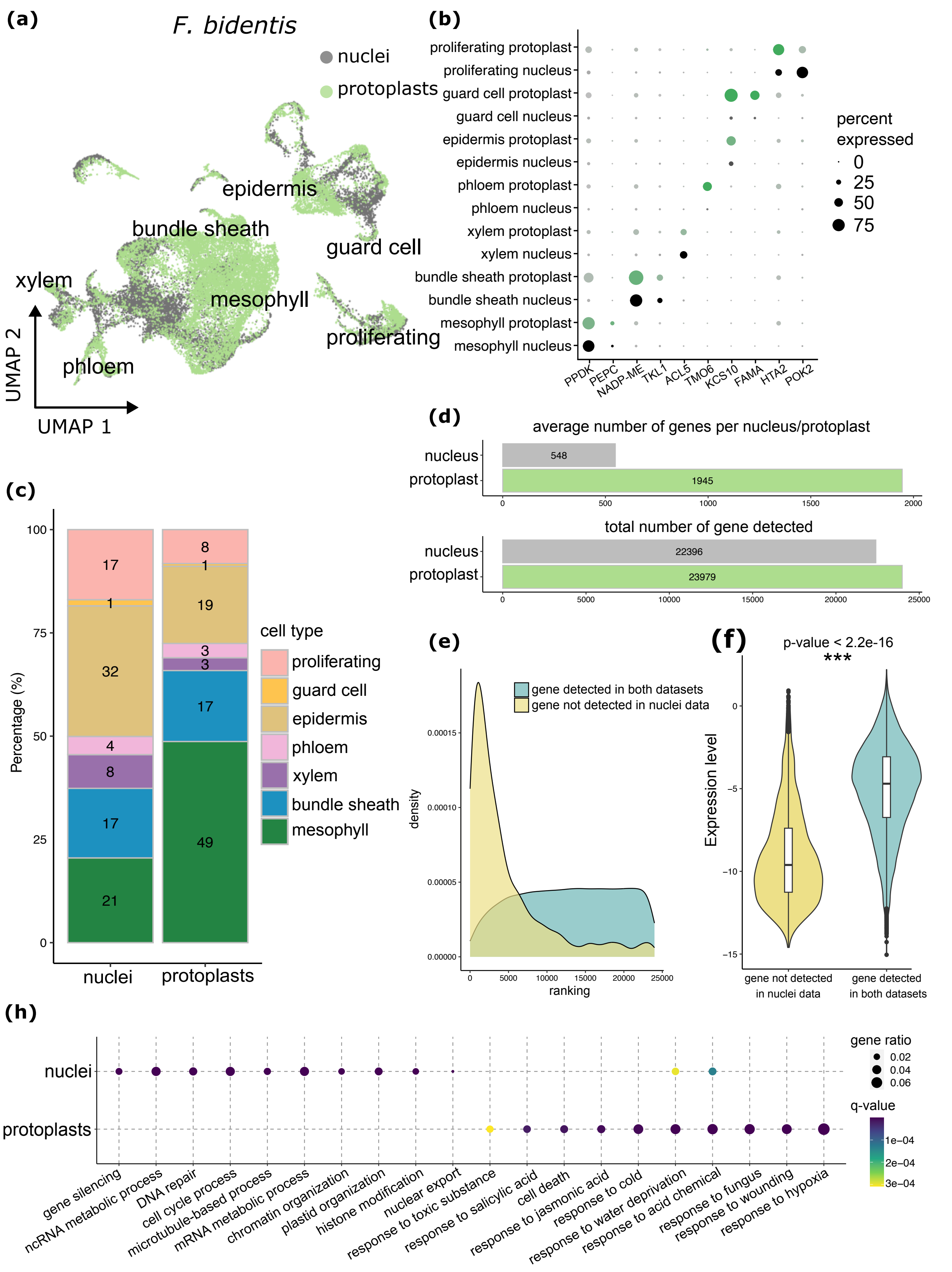


**Fig. S7**

Comparison of single-nucleus and single-protoplast transcriptome profiles in *F. bidentis* leaves. (a) UMAP visualization of combined protoplast and nuclei transcriptomic profiles. Major clusters annotated by cell-type. Nuclei shown as grey and protoplasts as green. (b) Dot plots showing the expression of marker genes for major cell types from either nuclei or protoplasts. CC/SE, companion cell/sieve element related cells. (c) Bar chart showing representation of each cell type estimated from nucleus or protoplast data based on expression of marker genes. CC/SE, companion cell/sieve element related cell. (d) Bar plot comparing the average number of genes per nucleus or protoplast, as well as the total number of genes detected in each profile. (e) Density plot showing the distribution of gene ranks, where yellow represents genes detected only in the protoplast profile, and teal genes detected in both nuclei and protoplast profiles. (f) Violin plot illustrating expression of genes detected in protoplasts only (yellow) or nuclei and protoplasts (teal). (h) Dot plots showing enriched Gene Ontology (GO) terms for genes that were differentially expressed between nuclei and protoplasts. metb., short for metabolic; proc., short for process.

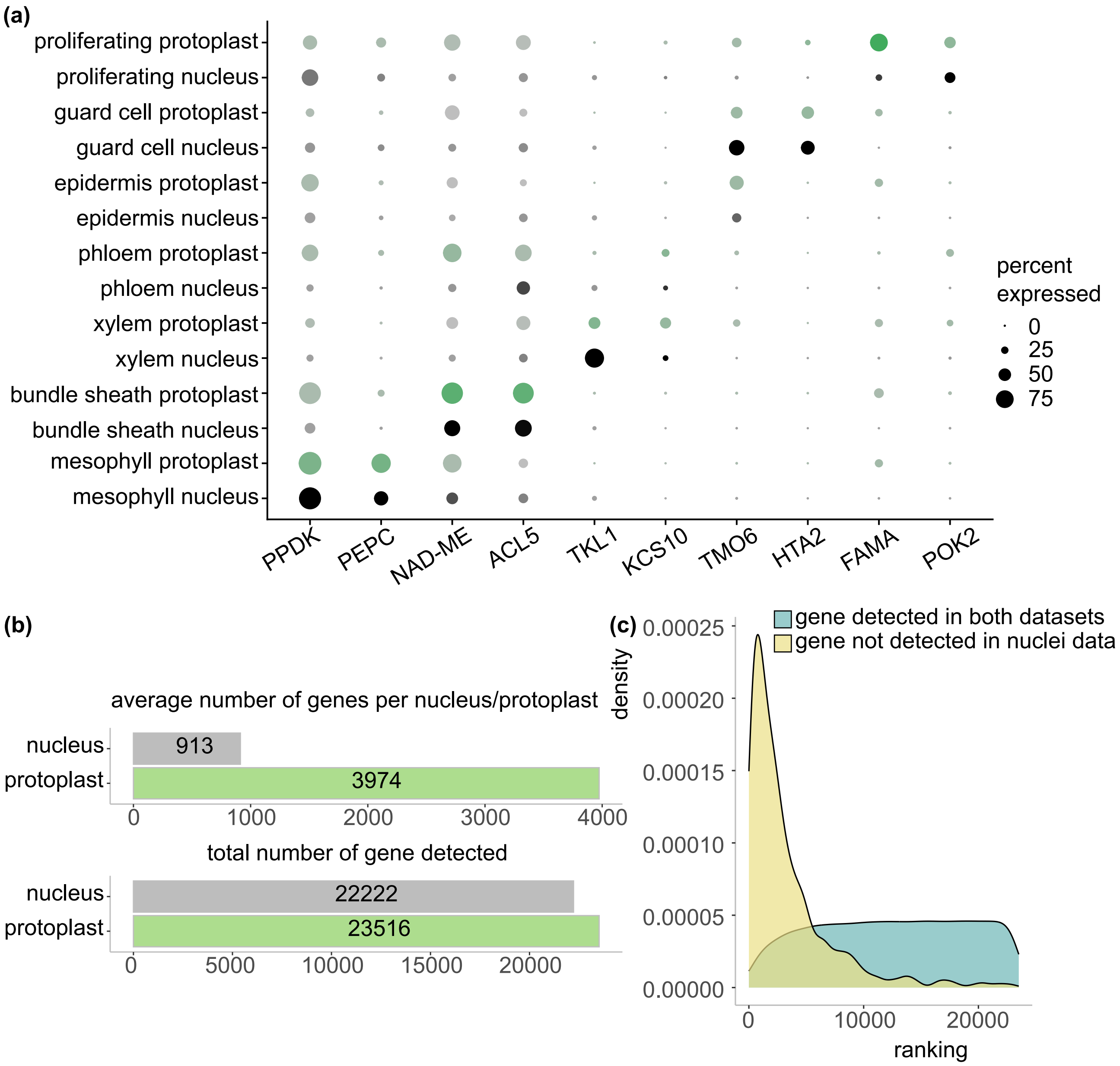


**Fig. S8**

Comparison of gene detection between single-nucleus and single-protoplast transcriptome profiles from *G. gynandra* leaves. (a) Dot plots showing the expression of marker genes for major cell types from either nuclei or protoplasts. CC/SE, companion cell/sieve element related cells. (b) Bar plot comparing the average number of genes per nucleus or protoplast, as well as the total number of genes detected in each profile. (c) Density plot showing the distribution of gene ranks, where yellow represents genes detected only in the protoplast profile, and teal represents genes detected in both nuclei and protoplast profiles.

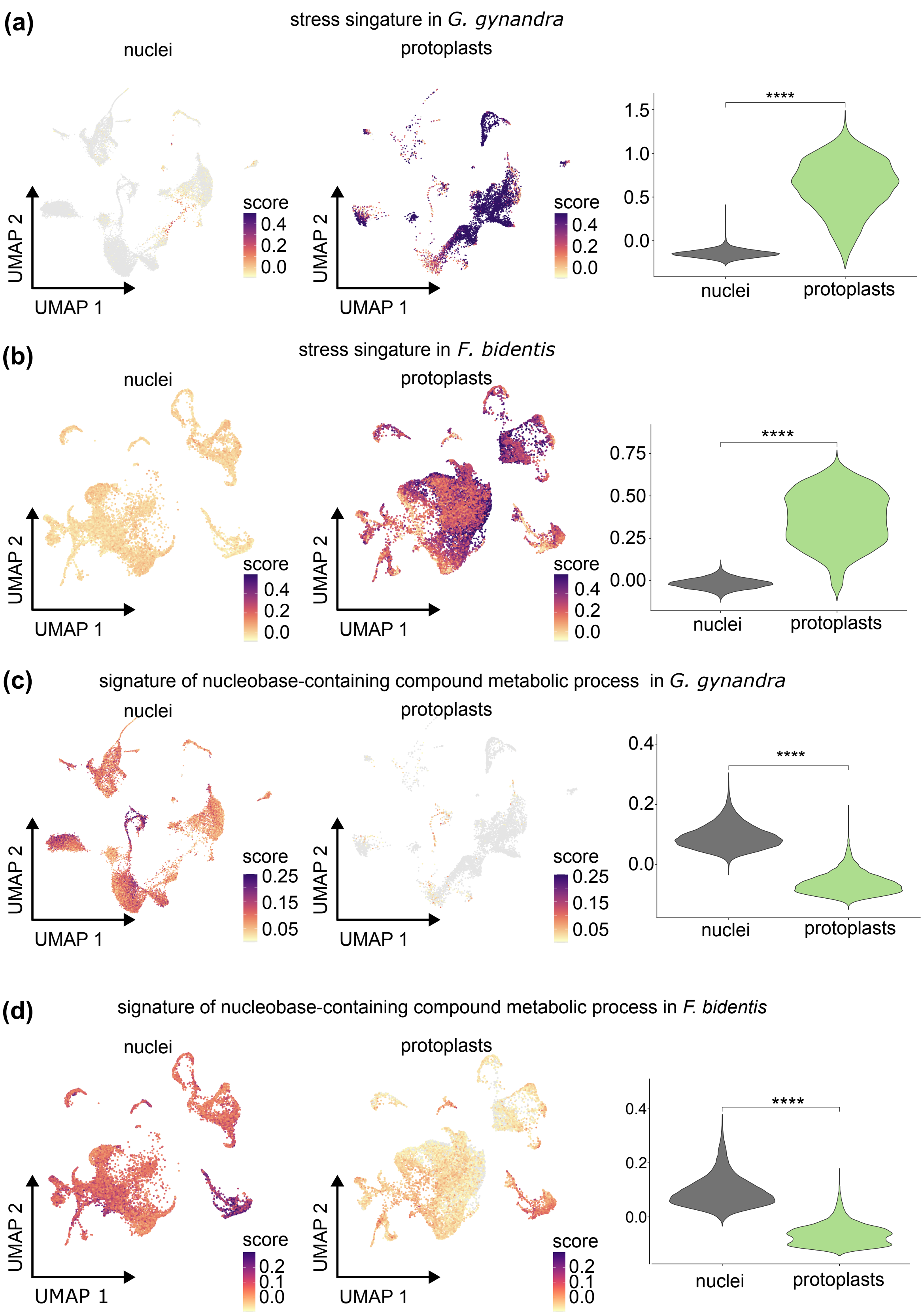


**Fig. S9**

Comparison of stress and nuclei signature scores. Expression of the stress score in nucleus and protoplast data of *G. gynandra* (a) and *F. bidentis* (b). Expression of the nucleus score in nucleus and protoplast data of *G. gynandra* (c) and *F. bidentis* (d). In each comparison, the first panel (left) shows the expression of the signature score in the single-nucleus transcriptome profile, the middle panel shows its expression in the single-protoplast transcriptomic profile, and the last panel (right) compares the signature score between the two datasets using a violin plot.

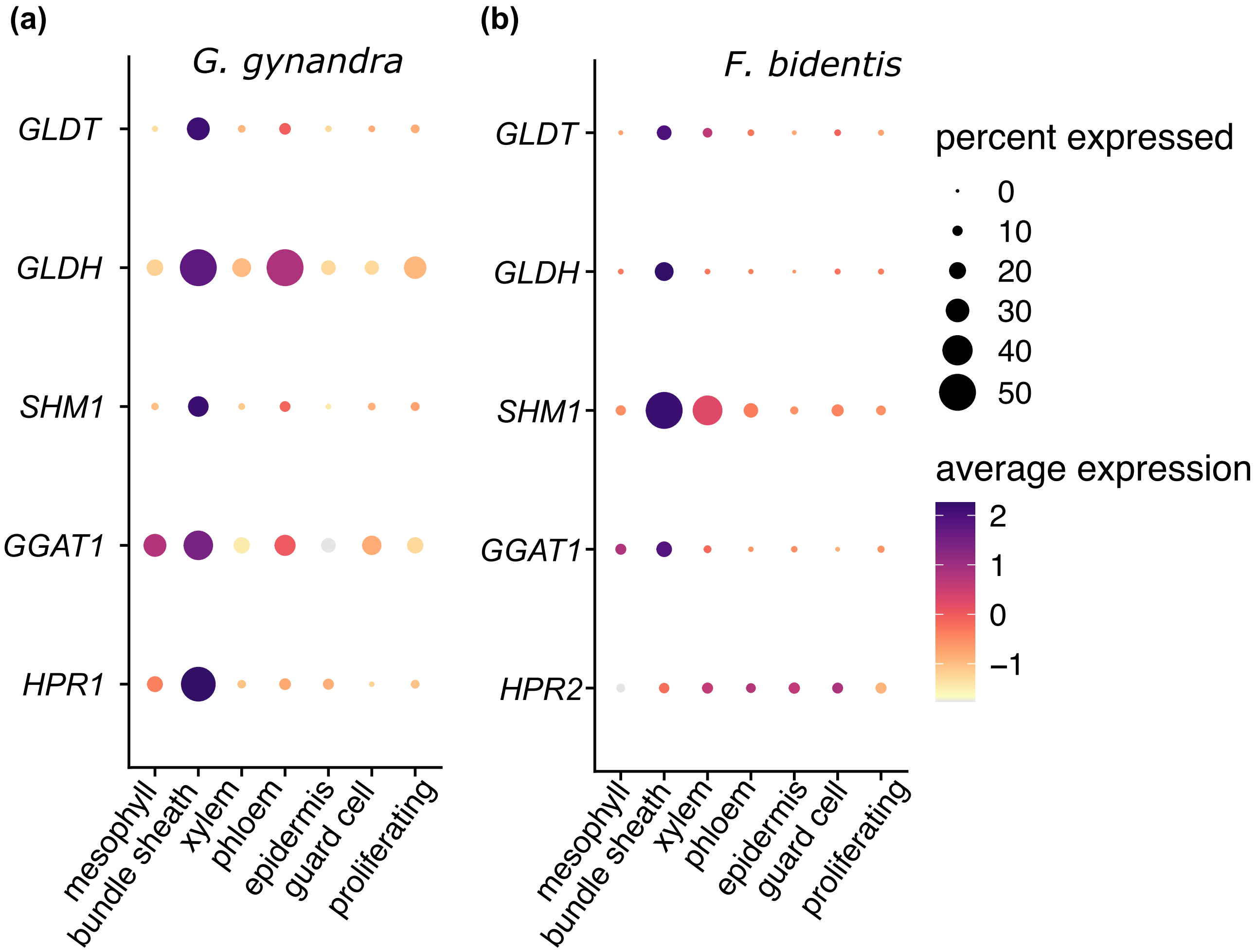


**Fig. S10**

Expression of photorespiration genes. Dot plots showing transcript abundance of photorespiration genes in different cell types of *G. gynandra* (a) and *F. bidentis* (b).

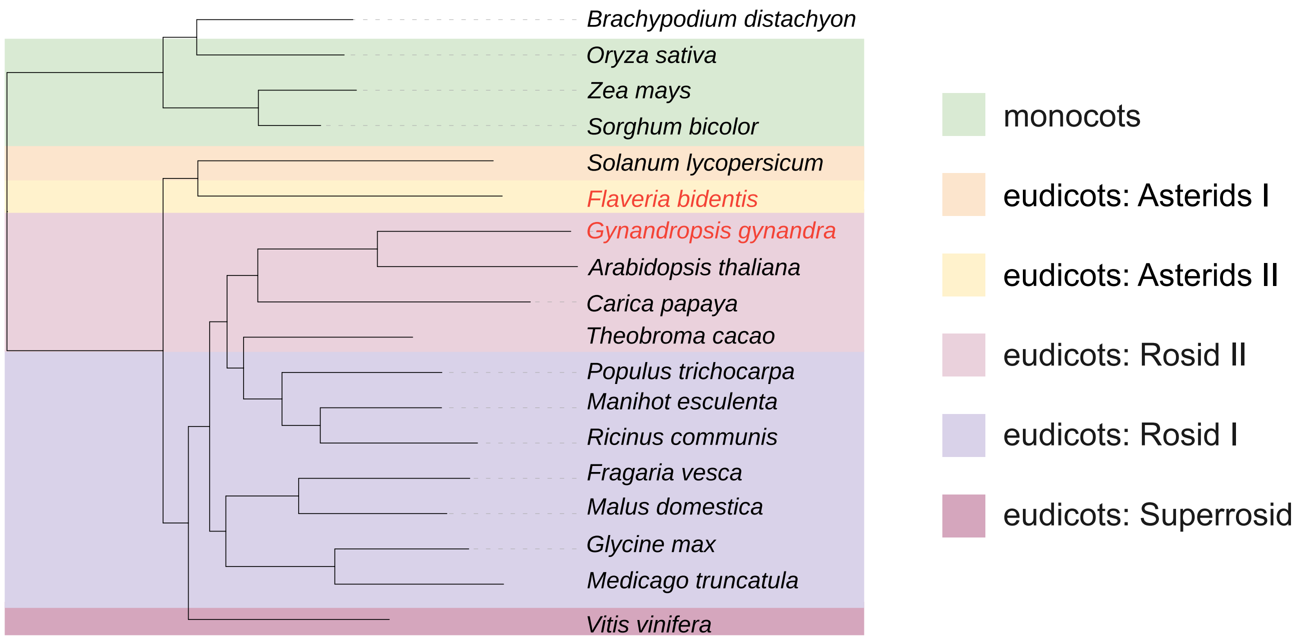


**Fig. S11**

Phylogenetic reconstruction of the species used to identify orthogroups.

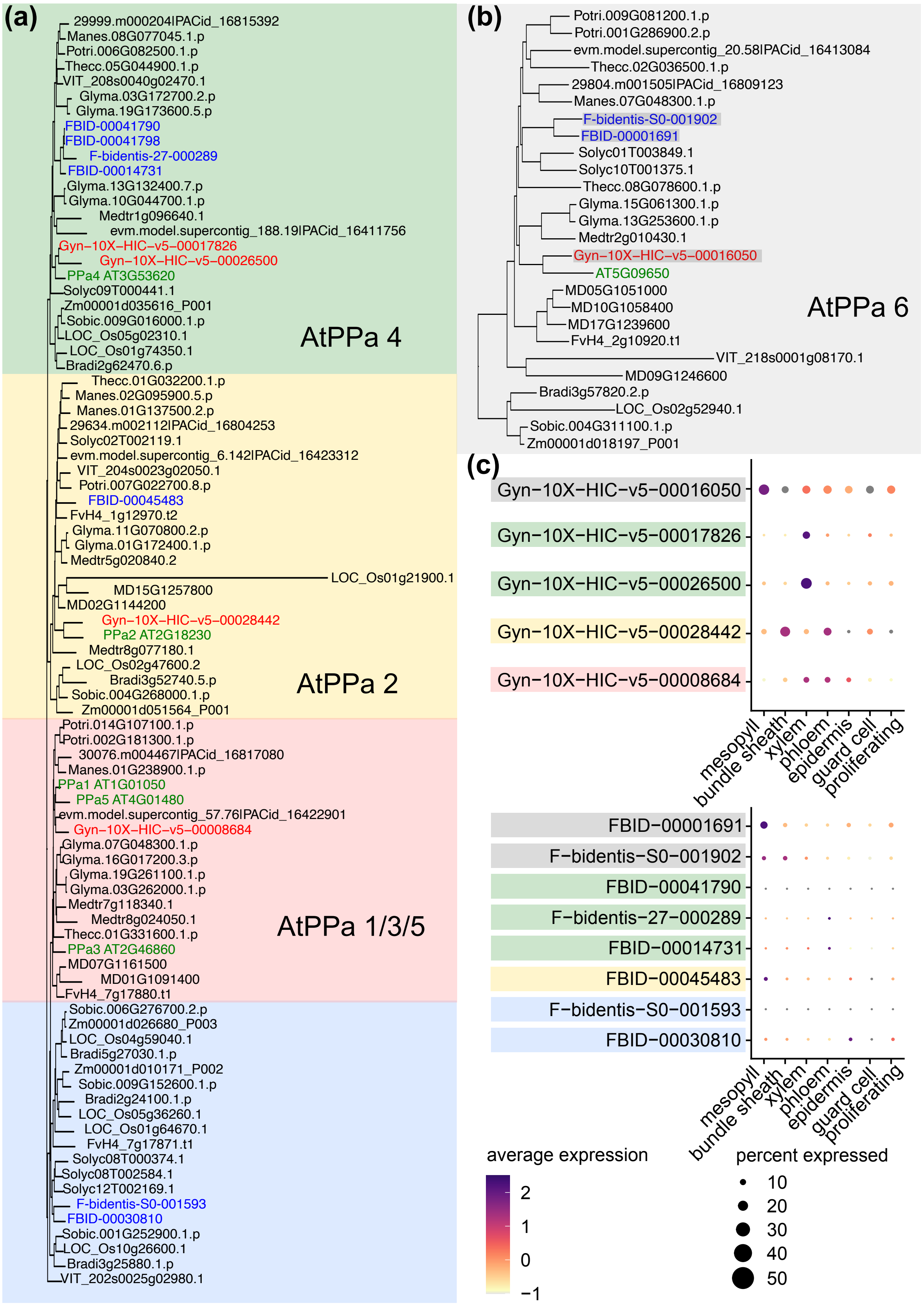


**Fig. S12**

Phylogenetic tree of the PPA orthogroups. (a) Phylogenetic tree of the AtPPA1-AtPPA5 orthogroup. (b) Phylogenetic tree of the AtPPA6 orthogroup. (c) Dot plots showing the expression of *PPA* genes for major cell types. *G. gynandra* and *F. bidentis* genes predicted to be chloroplast-localized are highlighted in the grey box.

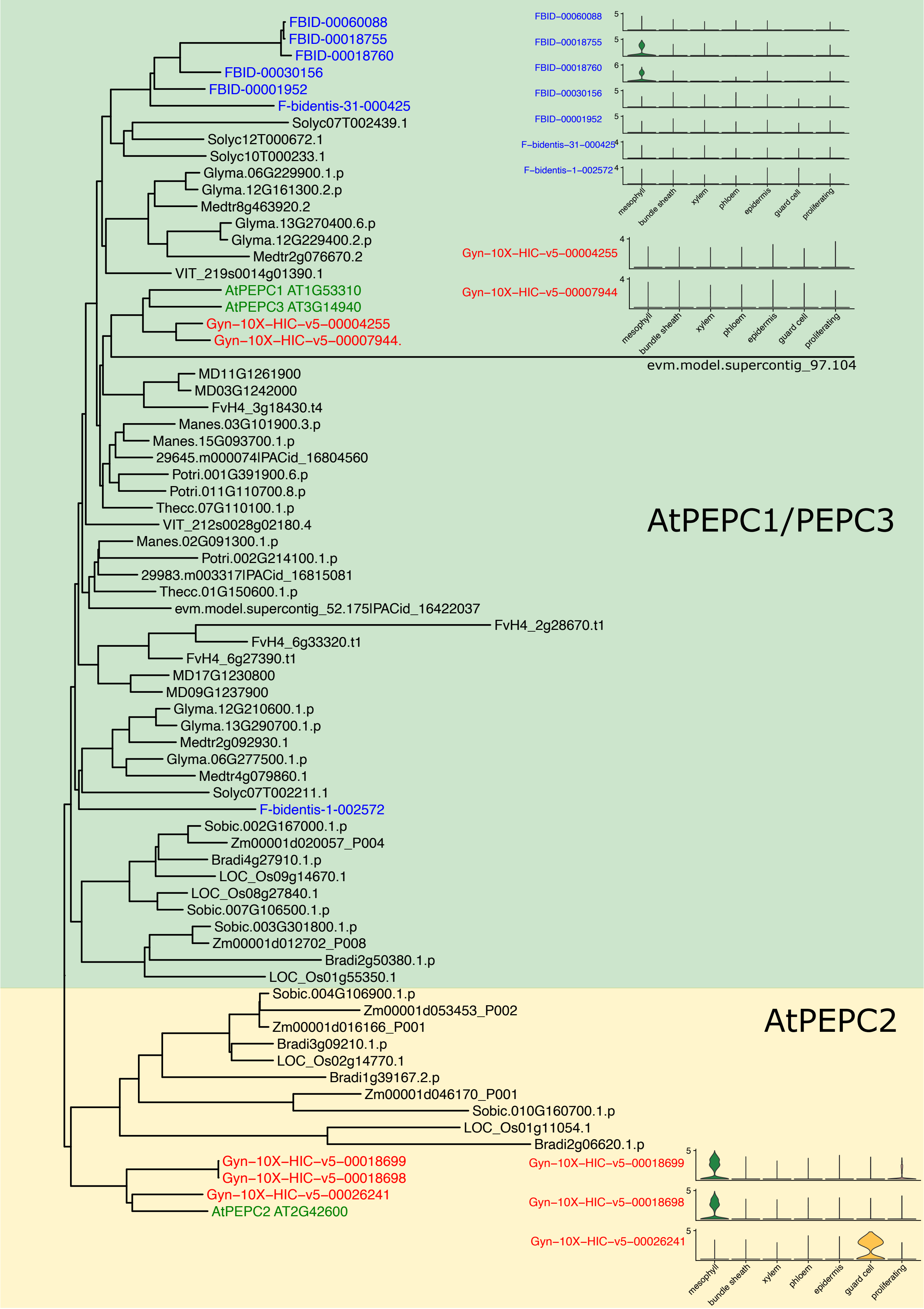


**Fig. S13**

Phylogenetic tree of the PEPC orthogroup. Expression of individual *PEPC* genes shown in violin plots.

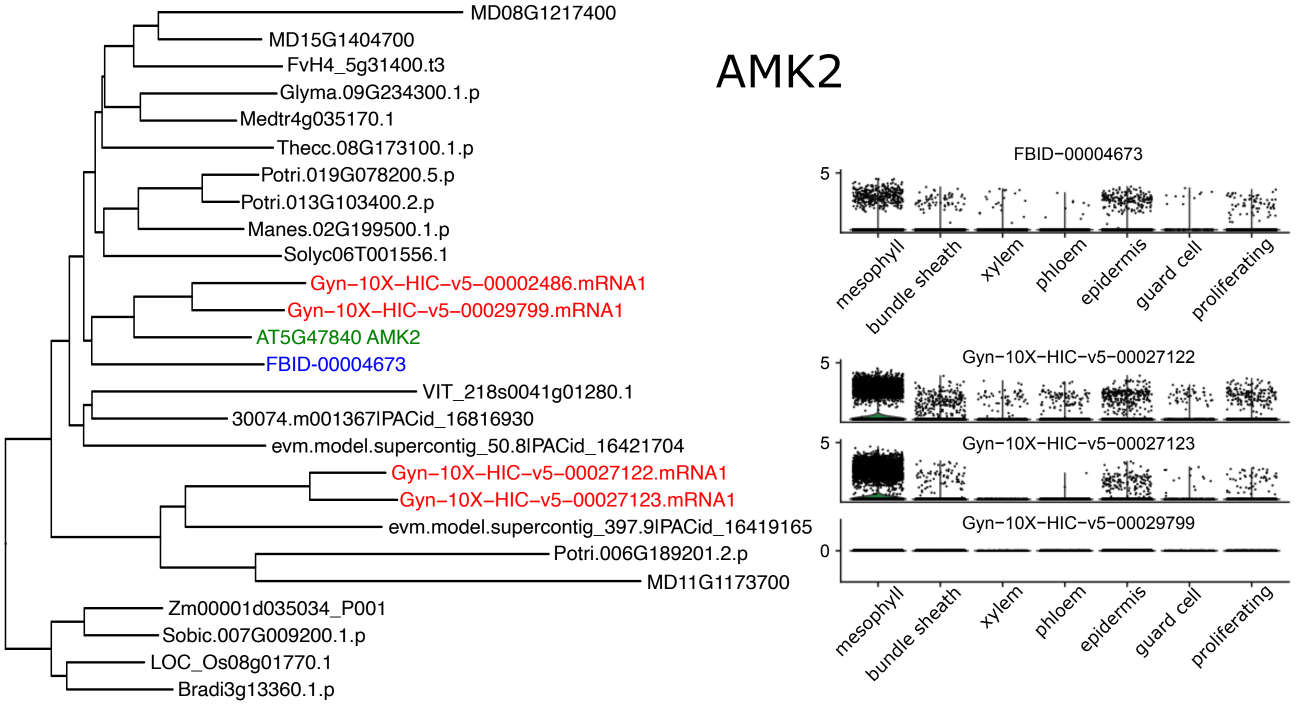


**Fig. S14**

Phylogenetic tree of the AMPK orthogroup. Expression of individual *AMPK* genes shown in violin plots.

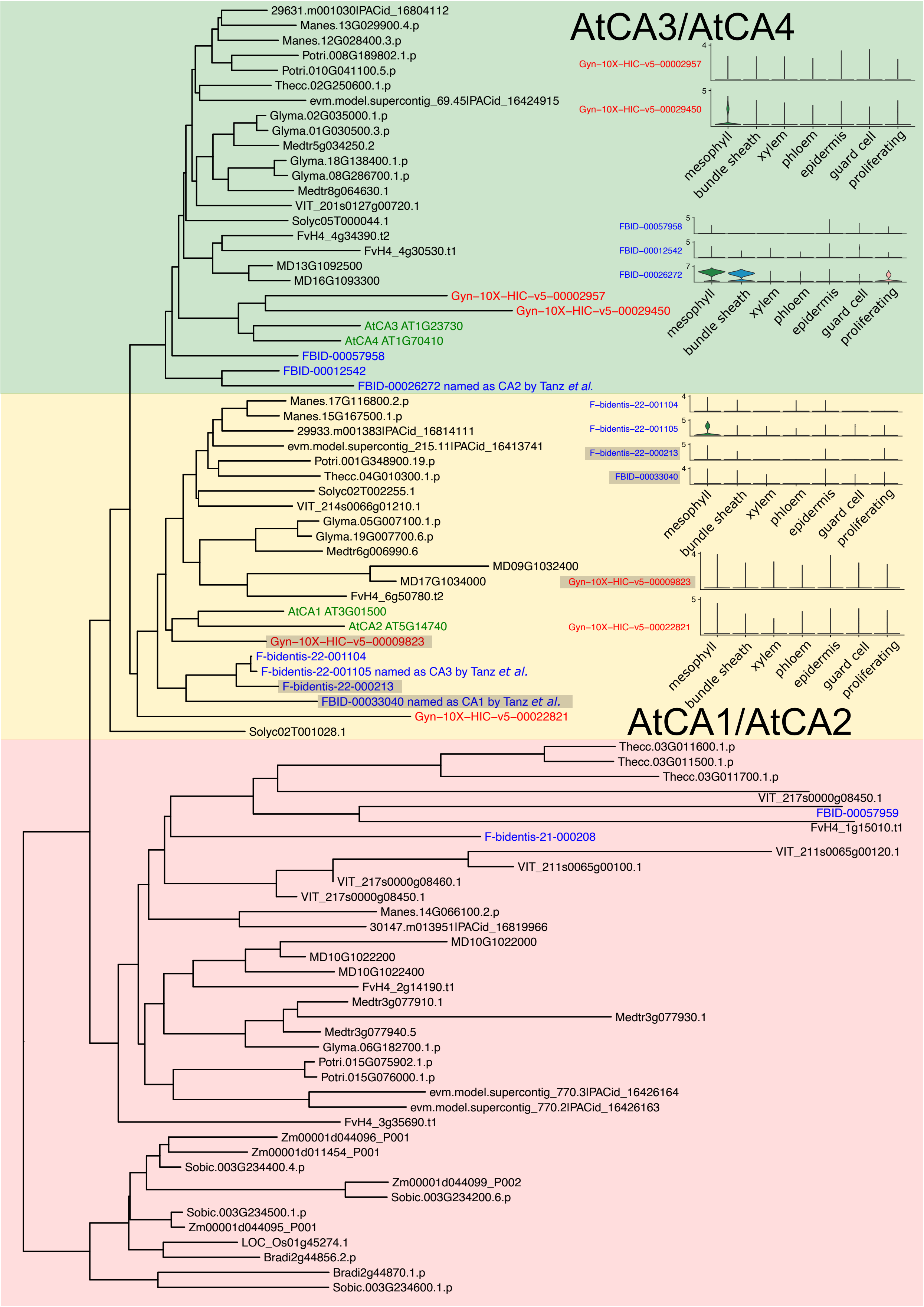


### Fig. S15

Phylogenetic tree of the CA orthogroup. Expression of individual *CA* genes shown in violin plots. *G. gynandra* and *F. bidentis* genes predicted to be chloroplast-localized are highlighted in the grey box.

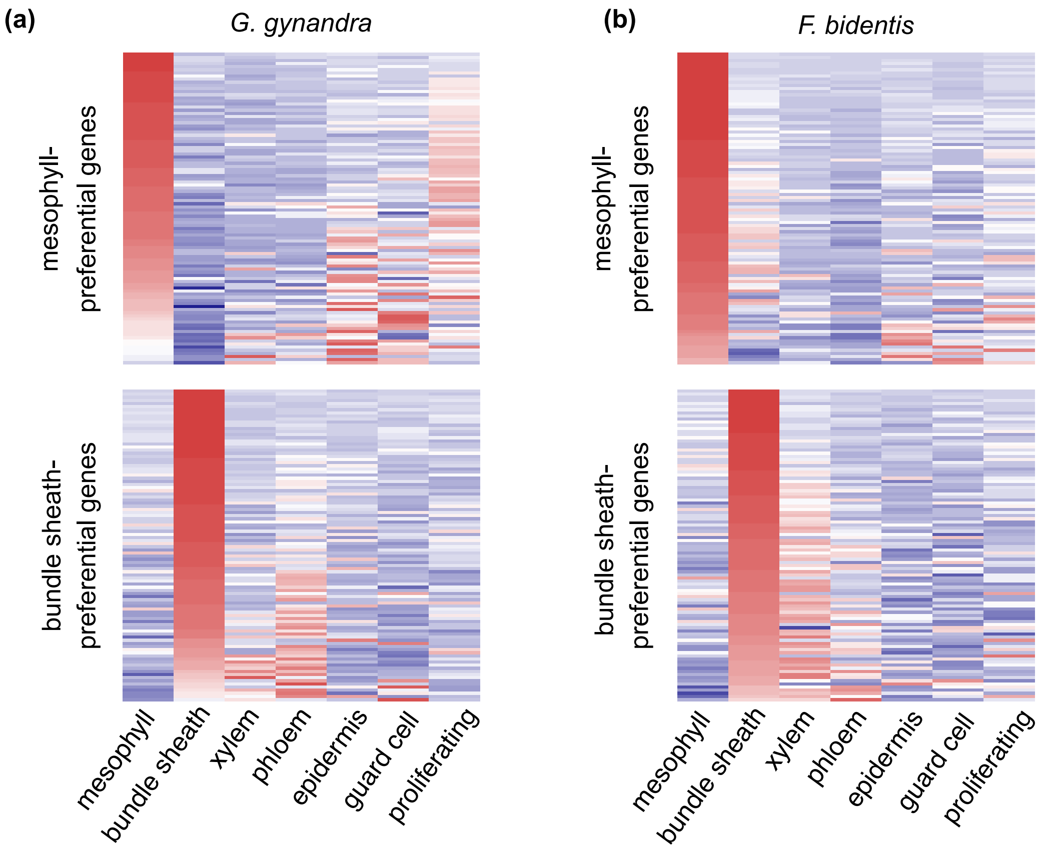


**Fig. S16**

Expression of differentially expressed genes between mesophyll and bundle sheath. Heatmap of the top 100 genes differentially expressed between mesophyll and bundle sheath cells of *G. gynandra* (a) and *F. bidentis* (b).

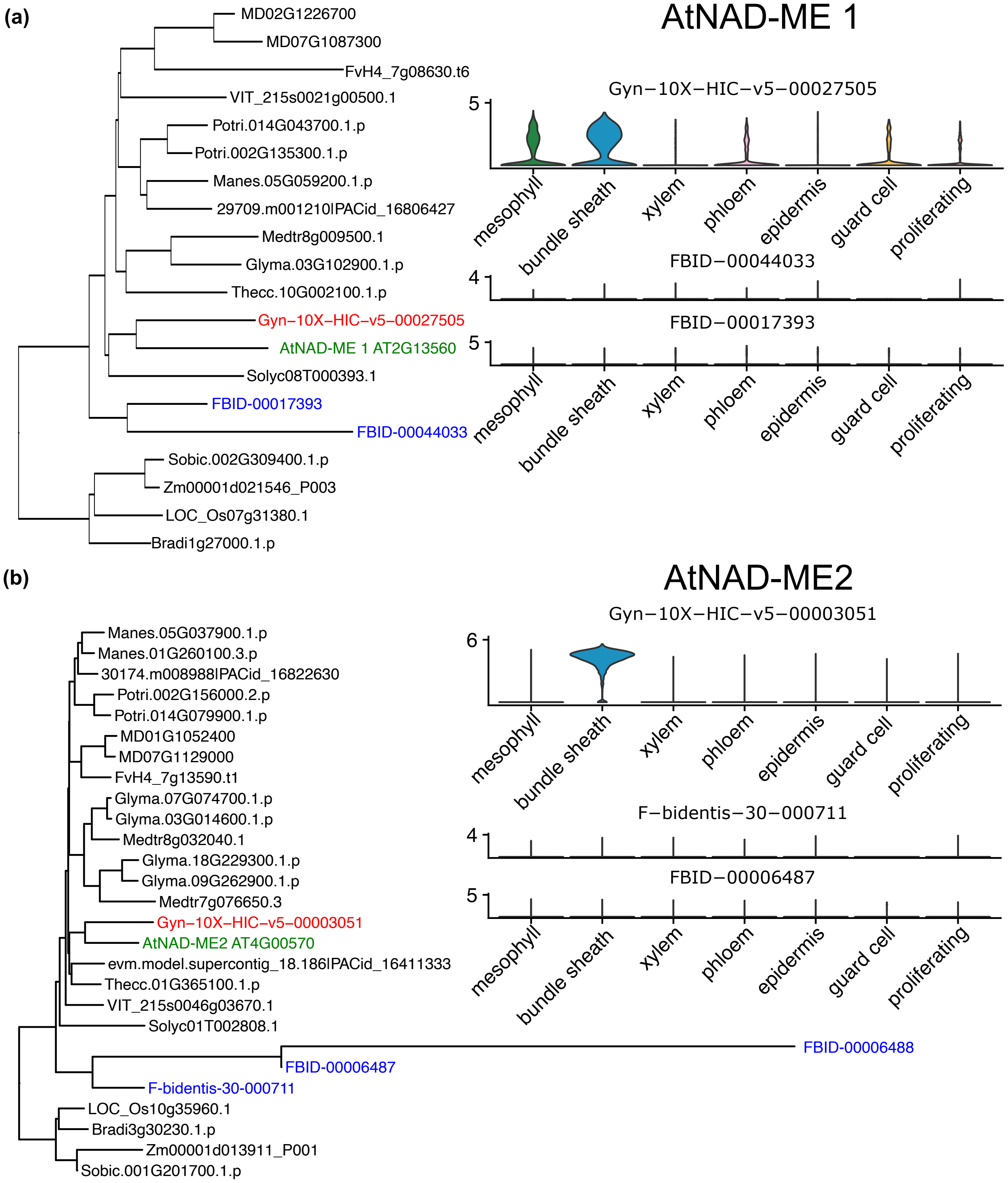


**Fig. S17**

Phylogenetic tree of the NAD-ME orthogroups. (a) Phylogenetic tree of AtNAD-ME1 (a) and AtNAD-ME2 orthologue (b), expression of individual *NAD-ME* genes shown in violin plots.

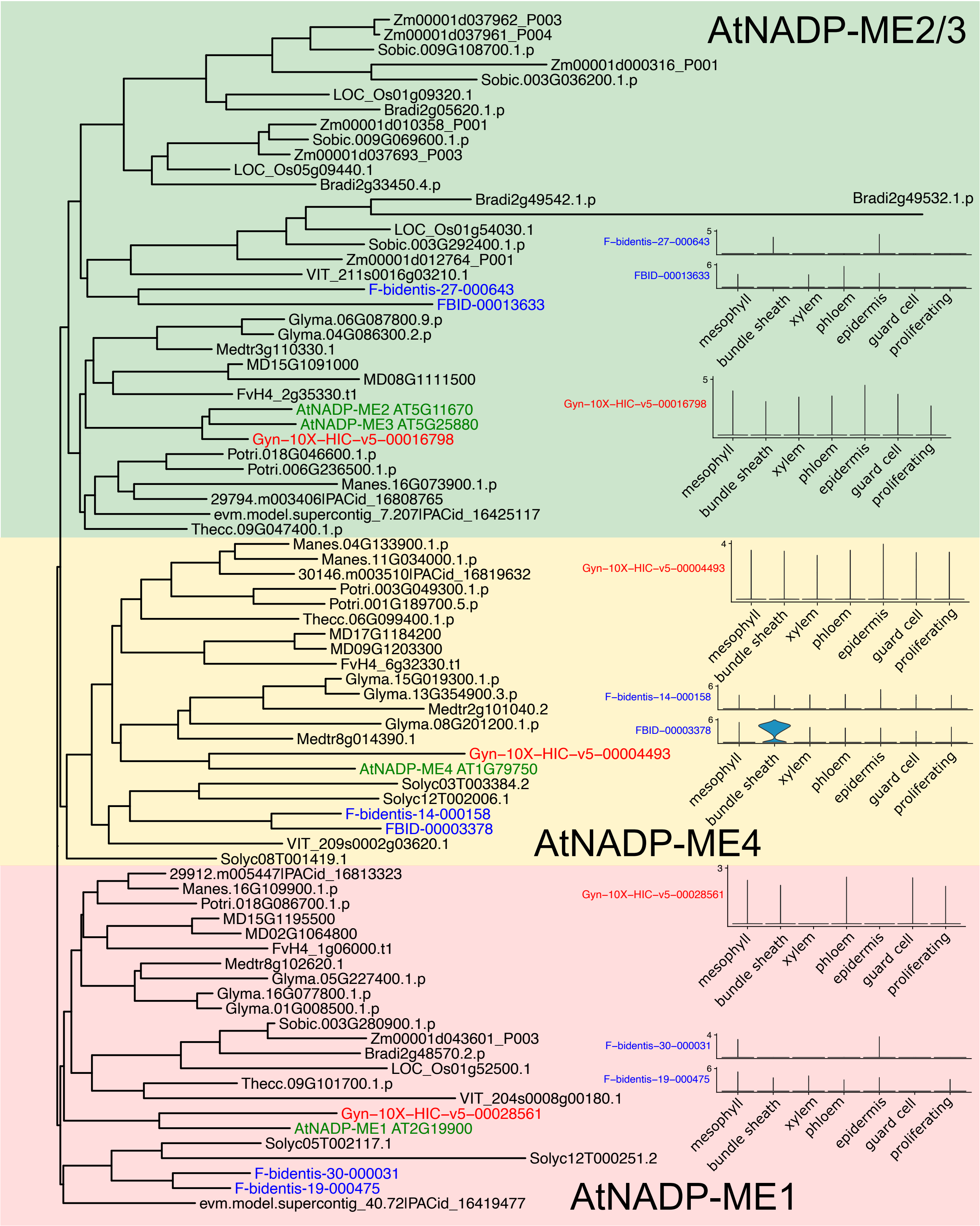


**Fig. S18**

Phylogenetic tree of the NADP-ME orthogroup. Expression of individual *NADP-ME* genes shown in violin plots.

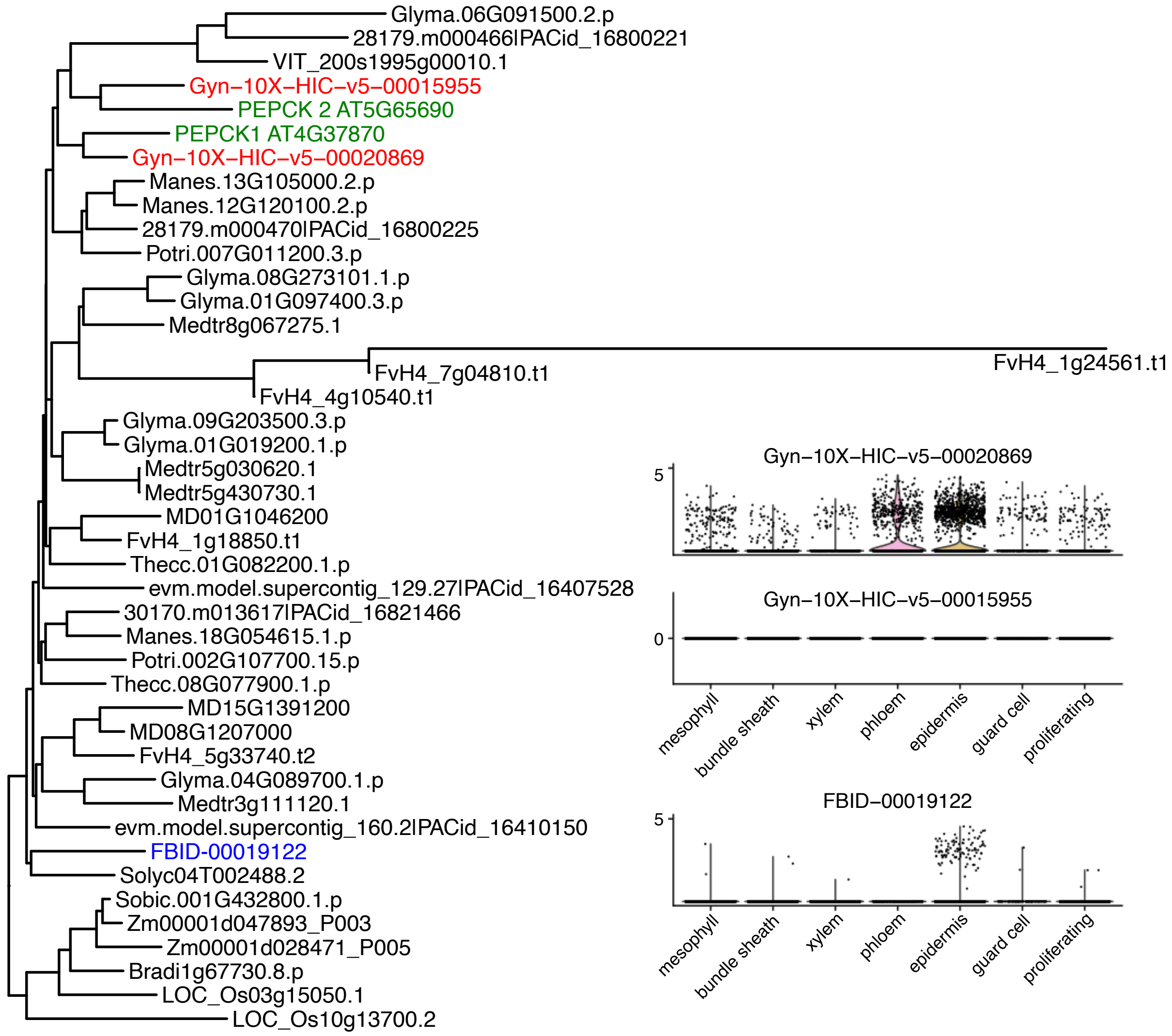


**Fig. S19**

Phylogenetic tree of the PEPCK orthogroup. Expression of individual *PEPCK* genes shown in violin plots.

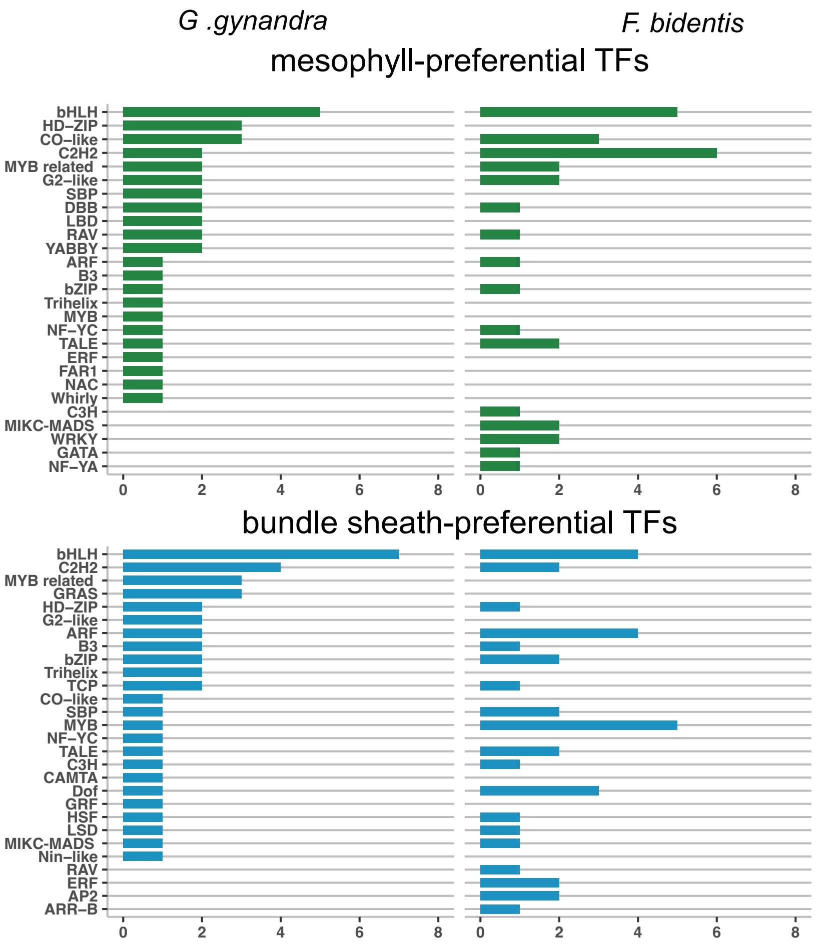


**Fig. S20**

Number of all mesophyll and bundle sheath preferential transcription factors characterised into different families in *G. gynandra* and *F. bidentis*.

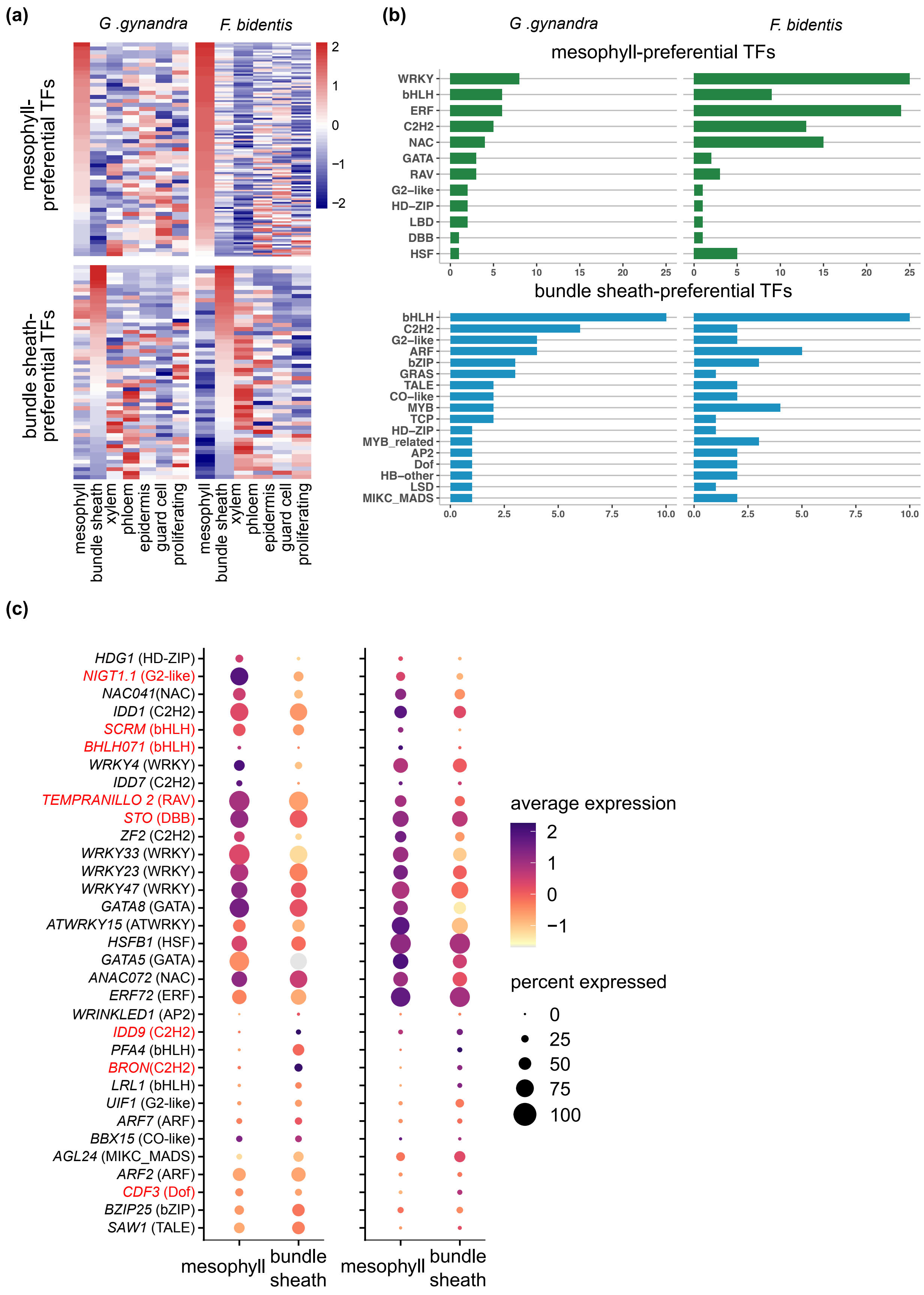


**Fig. S21**

Compartmentation of transcription factors between bundle sheath and mesophyll using protoplast data only. (a). Heatmap of genes differentially expressed between mesophyll and bundle sheath cells. (b) Number of common mesophyll and bundle sheath preferential transcription factors characterised into different families. (c) Transcript abundance of orthologous transcription factors that were preferentially expressed in either mesophyll or bundle sheath cells.
